# Phylodynamic insights on the early spread of the COVID-19 pandemic and the efficacy of intervention measures

**DOI:** 10.1101/2021.05.01.442286

**Authors:** Jiansi Gao, Michael R. May, Bruce Rannala, Brian R. Moore

## Abstract

We performed phylodynamic analyses of all available SARS-CoV-2 genomes from the early phase of the COVID-19 pandemic—combined with a novel dataset on contemporary global air-travel volume—to assess the efficacy of public-health measures on viral geographic spread. Globally, viral dispersal rates are significantly correlated with air-travel volume, and widespread international air-travel bans imposed against China by early February coincide with a significant reduction in geographic viral spread. In North America, the efficacy of this travel ban was temporary, possibly due to the lack of both containment measures against other infected regions and domestic mitigation measures. By contrast, in China, domestic mitigation measures were correlated with a long-term reduction in viral spread, despite repeated international introductions. Our study supports a role for both targeted international containment and domestic mitigation measures as critical components of a more comprehensive public-health strategy to mitigate future outbreaks caused by the emergence of novel SARS-CoV-2 variants.

**One sentence summary:** Phylodynamic analyses reveal that variation in rates of early geographic spread of COVID-19 are correlated with intervention measures.

---

The COVID-19 pandemic emerged in Wuhan, China, in late 2019, and rapidly established a global distribution by early March, 2020 *(1)*, despite the implementation of various intervention efforts to slow the spread of the causative SARS-CoV-2 virus *(2)*. This crucial early phase of the pandemic provides a unique opportunity to explore the dispersal dynamics that led to the worldwide establishment of the virus and the efficacy of public-health measures to mitigate the spread of COVID-19. We distinguish between two kinds of intervention measures: *containment* measures (such as international travel bans) intended to prevent the introduction of the virus to uninfected areas, and *domestic-mitigation* measures (such as shelter-in-place policies) intended to restrict the transmission of the virus within an infected area. These issues have typically been explored using epidemiological models that simulate the geographic spread of the disease using historical (rather than contemporary) air-travel data *(3, 4)*, where the passenger volume between each pair of areas serves as a proxy for the rate of viral dispersal among those areas.

Here, we explore the global establishment and mitigation of SARS-CoV-2 using phylodynamic methods; this explicitly probabilistic approach models the process of viral dispersal among a set of discrete geographic areas *(5)*. The observations include the viral genomic sequences, and the times and locations of viral sampling. These data are used to estimate the parameters of the phylodynamic model, which include the phylogeny of the viral samples, the global dispersal rate (average rate of dispersal among all geographic areas) and the pairwise dispersal rates (rate of dispersal across a particular dispersal route that connects a pair of areas). Existing phylodynamic models either assume that the geographic process was constant over the entire history of the virus, or that the process was constant within two or more time intervals, *i.e.*, a ‘piece-wise constant’ process where the global and/or pairwise dispersal rates vary among two or more pre-defined time intervals *(6–8)*. Importantly, inferences under phylodynamic models provide summaries of epidemiologically relevant factors, such as the number of viral dispersal events between areas, and the evidential support that each dispersal route was important to the spread of the virus (i.e., the ‘dispersal dynamics’).

Here, we report results of phylodynamic analyses of all publicly available SARS-CoV-2 genomic sequence data (2598 genomes as of September 22, 2020) sampled during this crucial early phase of the COVID-19 pandemic to: (1) explore the degree to which the dispersal process of geographic dispersal varied between time periods with different containment and mitigation measures (i.e., to assess whether the process was piecewise-constant in nature); (2) reveal variation in the global rate of viral dispersal within each distinct time interval; (3) understand changes in the dispersal routes that were important to the spread of the virus over the early phase of the pandemic, and; (4) assess the impact of widespread air-travel bans with China on the global dispersal rate and dispersal dynamics of the COVID-19 pandemic, focusing on the efficacy of travel restrictions in mitigating viral introductions into North America. Our phylodynamic study is unique in combining a contemporary dataset on global air-travel volume with a comprehensive sequence dataset to develop an integrative perspective on the early spread of the COVID-19 pandemic. Insights from the early phase of the pandemic have the potential to inform policies to mitigate the impacts of newly emerging variants of SARS-CoV-2 *(9–12)*.

## Rates of global viral dispersal and volume of global air travel are significantly correlated

The process of geographic spread during the early phase of the COVID-19 pandemic varies substantially over four distinct time intervals, exhibiting both increases and decreases through time (Fig. 1B, dark blue). The significant decrease in the global viral dispersal rate between the second and third interval (with a boundary at February 2) coincides with the initiation of international air-travel bans with China (imposed by 33 countries and nation states by this date). The virus achieved a global distribution by January 31 (with reported cases in 83% of the study areas by this date *(1)*); from this point onward, daily global viral dispersal rates—inferred under a more granular phylodynamic model (Fig. 1B, light blue)—are significantly correlated (*p* = 3.25e^−6^, Pearson’s *r* = 0.68) with independent information on daily global air-travel volume (Fig. 1B, dashed orange line). This correspondence supports the role of air travel as a primary dispersal vector for SARS-CoV-2, and explains the immediate efficacy of air-travel restrictions in limiting the global spread of COVID-19.

**Figure 1:**
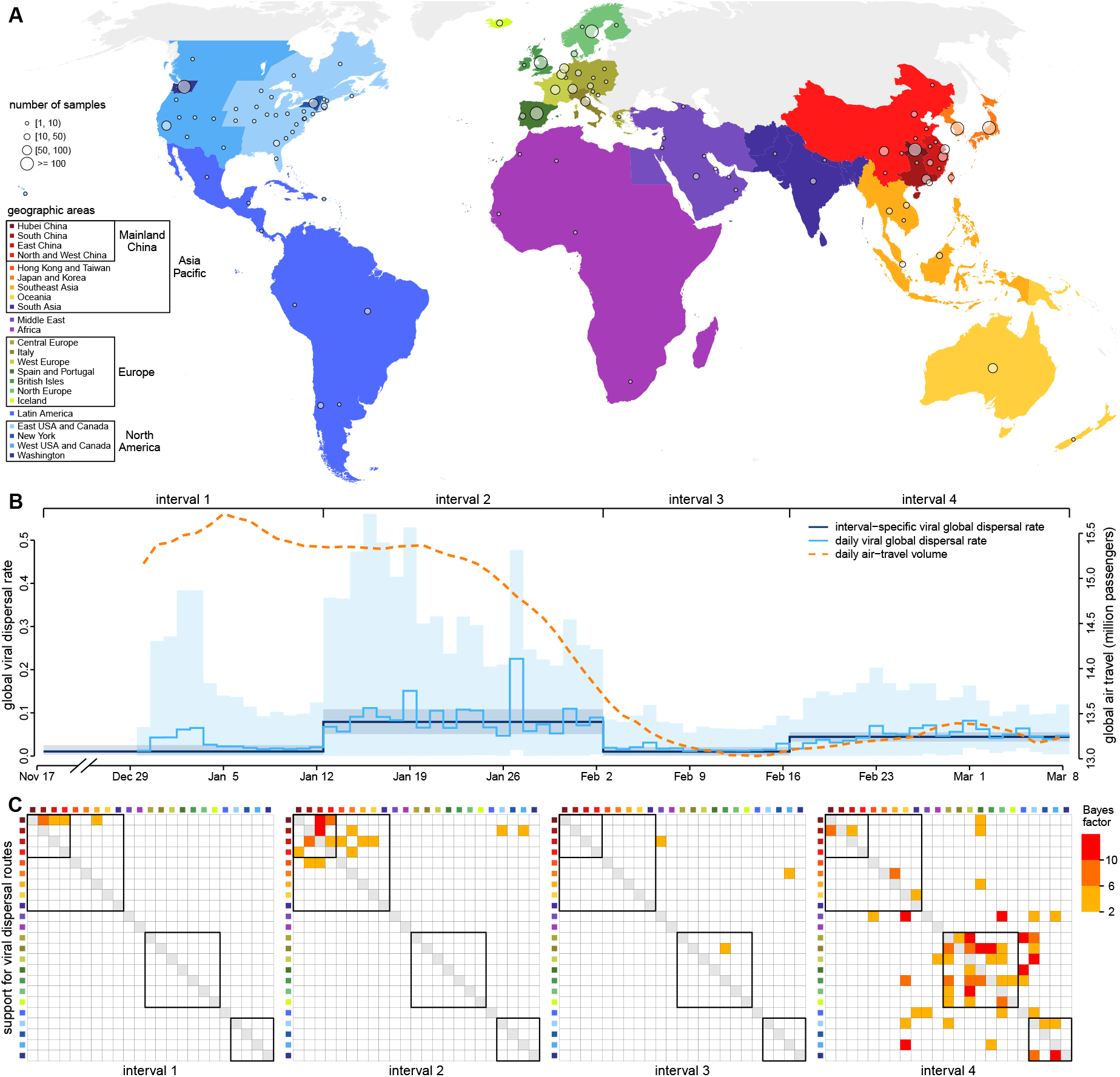
Phylodynamic analyses of SARS-CoV-2 genomes reveal temporal variation in rates and dynamics of viral dispersal during the early phase of the COVID-19 pandemic. **(A)**Our study includes a total of 2598 viral genomes collected between Dec. 24, 2019–Mar. 8, 2020 from 23 discrete geographic areas (colored regions); circles indicate the number and location of samples in our study. **(B)** The early phase of the COVID-19 pandemic is characterized by four time intervals spanning the MRCA of our sampled viruses (Nov. 17) to the end of our sampling period (Mar. 8). Each interval exhibits substantial differences in the global viral dispersal rate, i.e., the average rate of viral dispersal across all 23 areas (posterior mean [dark blue line], 95% credible interval [dark blue shaded area]). Notably, the global viral dispersal rate decreases sharply on Feb. 2, which coincides with the onset of international air-travel restrictions with China. More granular, daily estimates of the global viral dispersal rate (light blue) and independent data on global air-travel volume (the 7-day smoothed average number of passengers, dashed orange line) are significantly correlated. **(C)** The four time intervals also exhibit distinct dispersal dynamics. Heat maps indicate the level of evidential support (Bayes factors, inset legend, right) that a given dispersal route between a pair of geographic areas played a role in the spread of the virus. Boxes in each panel indicate groups of areas (inset legend, panel A). The first interval (Nov. 17–Jan. 12) is dominated by dispersal from Hubei to other areas in China, the second interval (Jan. 12–Feb. 2) exhibits more widespread dispersal within Asia and dispersal from China to North America, culminating in cosmopolitan dispersal in the fourth interval (Feb. 16–Mar. 8). The third interval (Feb. 2–Feb. 16)—immediately following the onset of international air-travel bans with China— exhibits a reduction in the number of viral dispersal routes, including disruption of the dispersal routes from China to North America.

## Viral dispersal dynamics vary over time and support the efficacy of air-travel restrictions

In addition to differences in the global viral dispersal rate, the four time intervals spanning the early phase of the COVID-19 pandemic also exhibit significant differences in viral dispersal dynamics, where the virus spread over characteristic dispersal routes within each time interval (Fig. 1C). As with the global viral dispersal rate, the number of dispersal routes by which the virus spread decreased substantially between the second and third interval (February 2), again providing evidence for the efficacy of the international air-travel bans in limiting the geographic progression of the disease. Importantly, our results provide significant support for direct viral dispersal from China to both eastern and western North America during the second interval; however, viral spread across these two dispersal routes declined significantly following the onset of the US air-travel ban with China (Fig. 1C). Within the third interval, we infer significant support for viral spread between only three pairs of geographic areas, where the corresponding dispersal routes were not impacted by air-travel bans, or where dispersal involved other means of travel. For example, our inference of viral spread across the dispersal route connecting Japan/Korea and western North America during this interval reflects passengers of the Diamond Princess cruise ship *(13)*.

## The impact of intervention measures on the number of viral dispersal events between areas

In addition to inferring dispersal rates and dispersal routes, our phylodynamic analyses also allow us to infer the number of viral dispersal events between pairs of areas, and thereby to assess the efficacy of intervention measures. Our results suggest that the early introduction of the virus into North America (prior to February 2) was dominated by dispersal events from China (Fig. 2A, orange). Later introductions of the virus into North America (after February 2) were increasingly dominated by dispersal events from outside China (Fig. 2A, purple). This explains the rapid but temporary effect of the air-travel ban with China: because the virus was concentrated in China prior to the onset of the air-travel ban, it appears to have provided an effective barrier to the major locus of viral infection. Nevertheless, the virus had already been seeded to many regions across the globe (80% of the study areas outside of China and North America, (1); *c.f.*, Fig. 2A, blue), which allowed increased dispersal of the virus into North America as the level of global infections rose. Moreover, once introduced, the virus proliferated within North America—as reflected by the increasing number of viral dispersal events within North America (Fig. 2A, green)— its spread unchecked by domestic travel restrictions or mitigation measures. China offers a striking contrast, both in terms of the strategies and impact of the measures it adopted to limit the spread of the COVID-19 pandemic. In contrast to targeted international air-travel bans, China imposed no restrictions on international travel, but instead adopted widespread domestic mitigation measures *(2, 14)*. This implementation was followed by a significant and sustained decrease in the number of viral dispersal events within China (Fig. 2B, green), despite persistent introductions from outside the country (Fig. 2B, purple).

**Figure 2:**
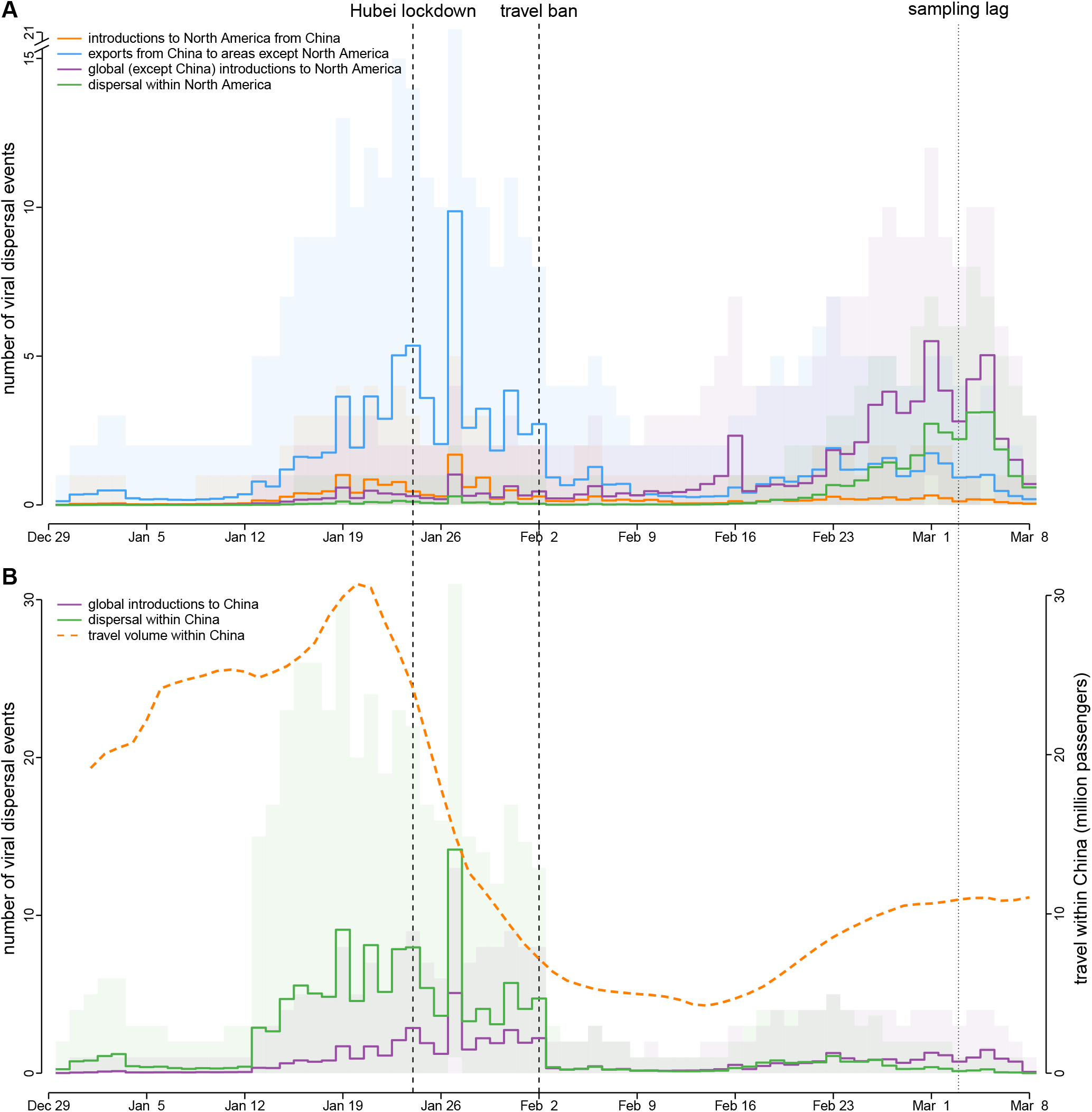
The impact of COVID-19 intervention measures adopted in North America and China. Each panel depicts the number of viral dispersal events between areas (posterior mean [solid lines], 95% credible interval [shaded areas]). **(A)** Prior to Feb. 2, China was the major source of viral introductions into North America (orange), which explains the efficacy of the air-travel ban imposed on China. Although the travel ban had a rapid effect, it was nevertheless temporary, as—by this time—many episodes of viral dispersal from China to other regions of the world had already occurred (blue). Accordingly, the virus continued to disperse into North America from regions outside of China (purple), and also to disperse among areas within North America (green). **(B)** The partial but temporary success of the North American containment measure contrasts with the long-term impact of mitigation strategies in China, which initiated a lockdown of the Hubei province on Jan. 24, and nation-wide travel restrictions by Jan. 26. These measures drastically reduced domestic travel within China (orange dashed line), resulting in a rapid, long-term reduction in the number of viral dispersal events within China (green), despite successive waves of international viral introductions (purple). Note that sampling lag causes the number of dispersal events close to the end of the sampling period to be underestimated.

## Assessing the robustness of phylodynamic inferences to sampling and methodological artifacts

The results of our phylodynamic analyses are based on a small and non-random sample of the total population of SARS-CoV-2 viruses that existed during the early phase of the COVID-19 pandemic. Nevertheless, our analyses include all currently available viral genomic sequences from this period, which are likely the vast majority of such data ever to be available (Fig. S3). Moreover, we performed a second series of analyses on a reduced dataset with fewer sequences (1271 samples available as of April 19) and different temporal and spatial sampling intensities (Fig. S4). Our results based on this reduced dataset are qualitatively identical to those based on the entire dataset (Figs. S9–S12), suggesting that our findings are robust to sampling artifacts.

Our results also depend on the assumed phylodynamic model, which inherently provides an imperfect description of the true process that gave rise to our data. To address concerns about the possible impacts of model misspecification, we first used Bayesian model-comparison methods to assess the relative fit of our SARS-CoV-2 datasets to a large pool of candidate phylodynamic models; these analyses indicate a decisive preference for our inference model (Table S4). We also used posterior-predictive simulation to assess the absolute fit of our inference model, which indicates that it adequately describes the process that gave rise to our SARS-CoV-2 datasets (Figs. S5 and S6). Finally, our exploration of a more granular phylodynamic model (which allows daily variation in the global dispersal rate) indicates that our inference model captures the major features of the underlying geographic process (Fig. 1B, dark and light blue).

## A phylodynamic perspective on the COVID-19 pandemic and intervention measures

Phylodynamic analyses of SARS-CoV-2 genomes provide a critical perspective that complements conventional epidemiological methods for exploring the geographic spread of the COVID-19 pandemic. For example, our finding of a significant correlation between global viral dispersal rates and global air-travel volume validates a key assumption of conventional epidemiological methods (that use air-travel volume as a proxy for viral dispersal rates between areas). Additionally, by providing an explicit historical perspective—reflected in the genealogical relationships among the viral samples—phylodynamic analyses provide the critical ability to tease apart the extent to which viral prevalence in a given area reflects within-area viral transmission *versus* between-area viral dispersal. The ability to make this distinction is essential for evaluating the relative efficacy of containment and mitigation measures on the spread of the virus.

Our phylodynamic analyses of the early phase of the COVID-19 pandemic demonstrate the potential of this approach to assess the absolute and relative efficacy of alternative containment and mitigation measures. Globally, targeted international air-travel bans with China coincided with a rapid and significant reduction in the global viral dispersal rate (Fig. 1B, blue), and both the number of dispersal routes (Fig. 1C) and the number of dispersal events from China (Fig. 2A, blue) to other areas. In North America, the air-travel ban with China is correlated with an abrupt disruption of viral dispersal routes (Fig. 1C) and viral dispersal events (Fig. 2A, orange) from China. However, our results also reveal the temporary efficacy of this targeted containment measure; in North America, the partial nature of this travel ban allowed continued dispersal of the virus into North America from areas outside of China (Fig. 2A, purple), and the lack of domestic mitigation measures allowed the virus to disperse within North America (Fig. 2A, green). By contrast, the domestic mitigation measures adopted by China appear to have provided a more effective long-term strategy, effecting a rapid and long-term disruption of domestic viral dispersal routes (Fig. 1C) and reducing the number of domestic dispersal events (Fig. 2B, green). Moreover, the number of domestic dispersal events within China (Fig. 2B, green) apparently remained low, despite continued waves of viral dispersal into the country (Fig. 2B, purple), effectively suppressing the re-establishment of the virus in China.

In summary, our results support the potential efficacy of both targeted-containment and domestic-mitigation measures, suggesting that both are integral components of a comprehensive public-health strategy to reduce the geographic spread of the COVID-19 pandemic. However, as the virus became more widespread, domestic mitigation measures became increasingly critical for the long-term control of the disease. Insights from analyses of the early dynamics of the COVID-19 pandemic have the potential to critically inform policies to limit the impact of possible future outbreaks of the disease associated with the emergence of novel variants of the SARS-CoV-2 virus.

## Author contributions

JG, MRM, BR, and BRM conceived the study, performed the analyses, made the figures, and wrote the manuscript. JG led data curation and computer code implementation. All authors read and approved the final manuscript.

## Data and materials availability

GISAID accession IDs of the SARS-CoV-2 sequences used in this study, as well as the flight-volume data (obtained from FlightAware, LLC) and intervention-measure data, are maintained in the GitHub repository (https://github.com/jsigao/covid19_phylodynamics_measure_efficacy_supparchive) and archived in the Dryad repository (https://datadryad.org/stash/share/qM_lKSgUqZ9jk3yKU5Fo5JurDb4iichKAnkt4PPh6Wg). The repositories also contain BEAST XML scripts used to perform the phylodynamic analyses, R scripts used to perform simulations and post processing, and a modified version of the BEAST program used for some of the analyses in this study.

## Supplementary materials

Materials and Methods (Section S0)

Supplementary Text (Sections S1–S3)

Figs. S1–S14

Tables S1–S8

References *(15–80)*

