## Supplementary Material for "Phylodynamic insights on the early spread of the COVID-19 pandemic and the efficacy of intervention measures"

#### Contents

**S0 Materials and Methods** **S2**
S0.1 Data Acquisition and Curation . . . . . S2
S0.2 Phylodynamic Analyses . . . . . S3
S0.3 Data and Code Availability . . . . . S5

**S1 Detailed Description of Data Acquisition and Curation** **S6**
S1.1 Epidemiological Data . . . . . S6
S1.2 Travel Data . . . . . S6
S1.3 Definition of Time Intervals . . . . . S8
S1.4 SARS-CoV-2 Genomic Sequence Data . . . . . S8

**S2 Detailed Description of Phylodynamic Analyses** **S11**
S2.1 Estimating a Dated Phylogeny for the Reduced SARS-CoV-2 Dataset . . . . . S11
S2.2 Evaluating Candidate Biogeographic Models . . . . . S14
S2.3 Joint Analyses of the Entire SARS-CoV-2 Dataset . . . . . S20
S2.4 Estimating Daily Global Viral Dispersal Rates . . . . . S24
S2.5 Assessing the Impact of Incomplete Viral Sampling . . . . . S27

**S3 Extending Phylodynamic Methods** **S33**
S3.1 Allowing both global dispersal rate and dispersal dynamics to vary under piecewise-constant model . . . . . S33
S3.2 Stochastic mapping of geographic histories under the piecewise-constant model . . . . . S34
S3.3 Posterior-predictive simulation for geographic models . . . . . S36

### S0 Materials and Methods

The results of our study are based on a complex and comprehensive series of computationally intensive analyses. In this section, we provide a high-level overview of our data collection and data analyses to clarify the rationale of our study, while directing readers to the corresponding subsections of the supplemental material that provide additional details on the various analyses that we performed.

#### S0.1 Data Acquisition and Curation

##### Epidemiological data

We used two types of epidemiological information in this study: (1) the number of confirmed COVID-19 cases, and; (2) the intervention measures involving China that were enacted during the early phase of the pandemic. We compiled a dataset of the number of confirmed COVID-19 cases recorded on each day for each country/province/state based on various sources (WHO 2020; DXY 2020; NHCPRC 2020; ECDC 2020; USCDC 2020) via two intermediate portals (Wu et al. 2020b; Dong et al. 2020; see Section S1.1 for details). We used these case-number data to assess the fraction of total cases represented by our genomic sequences, and to estimate the approximate date by which SARS-CoV-2 had spread to most geographic areas. We collected information on international travel bans with China and domestic mitigation measures within China from multiple sources (Wikipedia 2020a,b; Kraemer et al. 2020; Tian et al. 2020; Hsiang et al. 2020; Lai et al. 2020; see Section S1.1 for details).

##### Travel data

Our study incorporates three types of travel data: (1) global air-travel-volume data; (2) domestic travel-volume data within China, and; (3) domestic travel-volume data within the United States. We used the daily number of commercial passenger flights from FlightAware as a proxy for global air-travel-volume data (see Section S1.2 for details). We extracted domestic travel-volume data within China from the Baidu Migration dataset (Baidu Inc. 2020; Wang et al. 2014) (see Section S1.2 for details). We extracted domestic travel-volume data within the United States from two independent datasets release by Apple and Google, respectively (Apple Inc. 2020; Google LLC 2020) (see S1.2 for details).

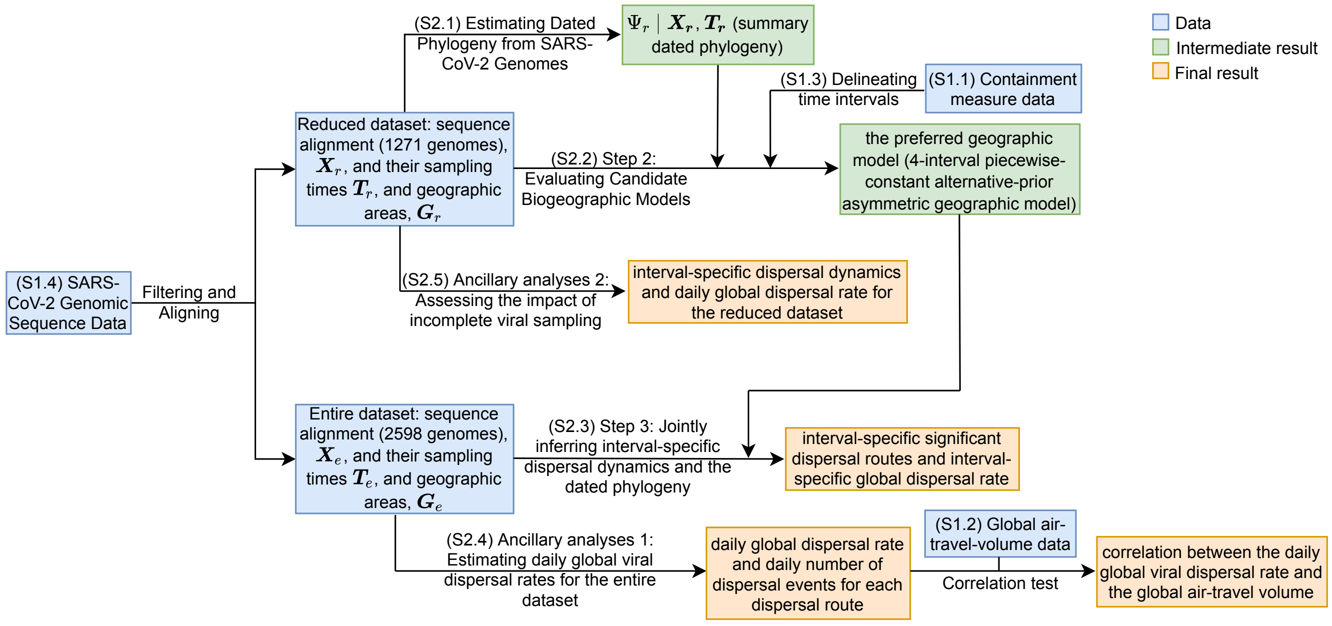


Figure S1: **Workflow of this study.**

##### Delineation of time intervals and geographic areas

To explore the dynamics of viral geographic dispersal in the early phase of the COVID-19 pandemic, we partitioned this phase into four time intervals of similar duration: (1) interval 1 from late 2019 (origin time) to January 12, 2020; (2) interval 2 from January 13, 2020 to February 2, 2020; (3) interval 3 from February 3, 2020 to February 16, 2020, and; (4) interval 4 from February 17, 2020 to March 8, 2020. Boundaries between these intervals coincide with the initiation of containment measures (*e.g.*, international travel bans with China) or other events associated with changes in the level of population movement (*e.g.*, start of the Spring Festival travel season); see Section S1.3 for details. For our phylodynamic analyses, we discretized the globe into geographic areas to study the early spread of SARS-CoV-2. We grouped geographically adjacent countries/territories (for non-focal regions) or states/provinces (for focal regions) to specify a total of 23 geographic areas (see Fig. 1A for the entire dataset and Fig. S11 for the reduced dataset).

##### SARS-CoV-2 genomic sequence data

We curated two genomic sequence datasets for this study, one with 1271 sequences (the "reduced dataset") and the other with 2598 sequences (the "entire dataset"). The reduced dataset was produced on April 19, based on all available SARS-CoV-2 genomic sequences from the Global Initiative on Sharing All Influenza Data (GISAID, Shu and McCauley 2017) as of that date. The entire dataset was produced by adding sequences that were available on GISAID as of September 22, 2020. See Section S1.4 for details on the sequence curation and alignment, and differences between the reduced and entire datasets.

#### S0.2 Phylodynamic Analyses

Our objective is to infer the joint posterior probability distribution of the viral phylogeny, divergence times, and biogeographic history under a composite phylodynamic model that is appropriate for the full SARS-CoV-2 dataset. The composite phylodynamic model is comprised of four main components: (1) a substitution model that describes the evolution of nucleotide sequences over the tree; (2) a branch-rate prior model that characterizes how rates of substitution vary across branches of the tree; (3) a branching-process model that specifies the prior distribution of the tree topologies and divergence times, and; (4) a biogeographic model that describes how viruses disperse between geographic areas.

For each of these components, there are numerous candidate models; the vast space of composite phylodynamic models makes it computationally prohibitive to evaluate the fit of each candidate model to the entire dataset. Accordingly, we adopt a three-step model-selection procedure: (1) we first estimate the dated phylogeny for the reduced dataset under a relaxed-clock model with biologically motivated specification of the substitution model, branch-rate prior model, and branching-process prior model; (2) we then use the resulting dated phylogeny to select among candidate biogeographic models, and; (3) finally, we performed joint inference of the dated phylogeny and biogeographic history for the entire dataset using the preferred composite phylodynamic model.

##### Step 1: Estimating the dated phylogeny of the reduced dataset

We inferred a dated phylogeny by performing Bayesian analyses of the reduced SARS-CoV-2 sequence dataset under a relaxed-clock model, which includes the first three of the four model components of the composite phylodynamic model: (1) a substitution model; (2) a branch-rate prior model, and; (3) a branching-process prior model.

Specifically, we specified a partitioned substitution model to accommodate possible variation in the evolutionary process across genomic regions. The SARS-CoV-2 genome is comprised of 11 gene regions (Table S1 lists these gene regions and their corresponding coordinates in the reference genome). We partitioned the SARS-CoV-2 genomes into six data subsets, with three subsets for the ORF1ab gene region (one for each codon position), and three subsets for the remaining ten combined gene regions (one for each codon position). For each of these data subsets, we specified an independent TN93 substitution model (Tamura and Nei 1993). We used partition-specific rate multipliers to capture differences in the substitution rate across the six data subsets. We specified a discrete-gamma model to accommodate substitution rate variation across sites within each data subset. Our preliminary analyses specified an independent discrete-gamma model for each of the six data subsets, which revealed a similar degree of among-site rate variation within each data subset (*i.e.*, with similar posterior estimates of the six $\alpha$-shape parameters). Accordingly, to decrease model complexity (and reduce MCMC issues), we specified a shared, discrete-gamma model (Yang 1994) to accommodate substitution-rate variation across sites of the entire alignment. We specified an uncorrelated lognormal (UCLN) branch-rate prior model (Drummond et al. 2006; Li and Drummond 2012; Rannala and Yang 2007) by drawing i.i.d. rate multipliers for each branch from a shared underlying lognormal distribution, where the parameters of this distribution (mean and standard deviation) are estimated from the data. For the branching-process prior model, we used a coalescent model with exponential population growth.

We performed MCMC simulations to approximate the joint posterior distribution of the relaxed-clock model parameters and the dated phylogeny using `BEAST` (Suchard et al. 2018). We then used `TreeAnnotator` to generate a summary phylogeny from the combined posterior sample of dated phylogenies as a maximum clade credibility (MCC) tree. We provide a more detailed description of these analyses in Section S2.1. The phylogeny inferred from these analyses was used in the next step to evaluate candidate biogeographic models.

##### Step 2: Evaluating candidate biogeographic models using the reduced dataset

Biogeographic models describe the history of viral dispersal over the dated phylogeny as a continuous-time Markov chain (CTMC). For a biogeographic history with $k$ discrete areas, a constant biogeographic model is fully specified by a $k \times k$ instantaneous-rate matrix, $\mathbf{Q}$, where an element of the matrix, $q_{ij}$, is the instantaneous dispersal rate from area $i$ to area $j$. This element is specified as $q_{ij} = r_{ij}\delta_{ij}$, where $r_{ij}$ is the relative rate of dispersal from $i$ to $j$, and $\delta_{ij}$ indicates whether the dispersal route from $i$ to $j$ exists ($\delta_{ij} = 1$) or not ($\delta_{ij} = 0$). More complex, piecewise-constant biogeographic models allow the average dispersal rate and/or the dispersal dynamics (*i.e.*, the instantaneous-rate matrix) to differ between two or more pre-specified intervals of the geographic history.

We explored a pool of candidate biogeographic models corresponding to possible combinations of: (1) symmetric and asymmetric $\mathbf{Q}$ matrices; (2) default and alternative priors on the total number of dispersal routes; (3) default and alternative priors on the average dispersal rate, and; (4) models where the $\mathbf{Q}$ matrices and average dispersal rates are piecewise constant over 1, 2 (with a boundary at February 2, 2020), or 4 (with boundaries at January 12, February 2, and February 16, 2020) pre-specified intervals.

We assessed the *relative fit* of these candidate biogeographic models to our reduced SARS-CoV-2 dataset by computing Bayes factors using the marginal likelihood estimated for each model. We ran power-posterior MCMC simulations using `BEAST` (Suchard et al. 2018), and then computed marginal likelihoods using both thermodynamic-integration (Lartillot and Philippe 2006) and stepping-stone (Xie et al. 2011; Baele et al. 2012) estimators.

We also assessed the *absolute fit* of each candidate biogeographic model to our reduced SARS-CoV-2 biogeographic dataset using posterior-predictive simulation (Gelman et al. 1996). For each model, we first inferred the joint posterior distribution from the observed biogeographic data (*i.e.*, the geographic location of each of the sequences in our reduced SARS-CoV-2 dataset) by performing MCMC simulations using `BEAST` (Suchard et al. 2018). We then simulated predictive datasets by repeatedly sampling at random from the corresponding joint posterior probability distribution for the given the model. Finally we generated posterior-predictive distributions from each predictive dataset under various test statistics, which measure the discrepancy between the observed dataset and the simulated dataset. We provide a more detailed description of these analyses in Section S2.2. The preferred biogeographic model identified by these analyses was used in our subsequent joint phylodynamic analyses, described below.

##### Step 3: Joint phylodynamic inference of the entire dataset

We performed joint inference of the phylogeny, divergence times, and biogeographic history using the entire SARS-CoV-2 dataset based on a phylodynamic model that includes (1) a relaxed-clock model, and (2) a biogeographic model. The relaxed-clock model specified in these joint analyses is identical to that specified in Step 1 (with minor changes in the prior specification to accommodate differences in viral sampling). The biogeographic model specified in these joint analyses is identical to the biogeographic model selected in Step 2: the 4-interval piecewise-constant alternative-prior asymmetric geographic model.

We performed MCMC simulations to approximate the joint posterior distribution of the phylodynamic-model parameters from the entire SARS-CoV-2 dataset using `BEAST` (Suchard et al. 2018). We also performed posterior-predictive simulations to confirm that the preferred biogeographic model provides an adequate fit to the entire SARS-CoV-2 dataset under the joint inference. We provide a more detailed description of these analyses in Section S2.3.

##### Ancillary analyses 1: Estimating daily global viral dispersal rates for the entire dataset

We performed additional analyses to explore the correlation between daily global air-travel volume and global SARS-CoV-2 dispersal rates during the early phase of the COVID-19 pandemic. We first estimated the daily global dispersal rate using the entire SARS-CoV-2 dataset under a more granular phylodynamic model that allows the average dispersal rate to vary from day to day. We then computed the correlation between these estimates and independent information on the daily volume of global air travel during this period. The phylodynamic model we specified for these analyses was identical to that used in Step 3 (joint phylodynamic inference), except that we further discretized the time intervals for the global dispersal rate to vary among days. In these analyses, we accommodated phylogenetic uncertainty by averaging over the marginal posterior probability distribution of dated phylogenies inferred in Step 3. We performed MCMC simulations to approximate the joint posterior distribution of the daily-rate model parameters using `BEAST` (Suchard et al. 2018). We then performed standard correlation test between the daily global air-travel volume and the estimated mean daily global SARS-CoV-2 dispersal rates by computing Pearson's r and the corresponding p-value, focussing on the period spanning from January 31 (by which date the virus achieved a global distribution) to March 8, 2020. We provide a more detailed description of these analyses in Section S2.4.

##### Ancillary analyses 2: Assessing the impact of incomplete viral sampling

We also performed additional series of analyses to assess the sensitivity of our results to incomplete and non-random sampling of SARS-CoV-2 sequences. Specifically, we replicated the entire series of analyses that we performed on the entire SARS-CoV-2 dataset—including joint inference of phylogeny, divergence times and biogeographic history, as well as analyses to infer daily global viral dispersal rates—for the reduced SARS-CoV-2 dataset, which has fewer (1271 vs 2598) sequences and different temporal and spatial sampling intensities. See Section S2.5 for details about these sensitivity analyses.

#### S0.3 Data and Code Availability

GISAID accession IDs of the SARS-CoV-2 sequences used in this study, as well as the flight-volume data (obtained from FightAware, LLC) and intervention-measure data, are maintained in the GitHub repository (`https://github.com/jsigao/covid19_phylodynamics_measure_efficacy_supparchive`) and archived in the Dryad repository (`https://datadryad.org/stash/share/qM_lKSgUqZ9jk3yKU5Fo5JurDb4iichKAnkt4PPh6Wg`). The repositories also contain `BEAST` XML scripts used to perform the phylodynamic analyses, `R` scripts used to perform simulations and post processing, and a modified version of the `BEAST` program used for some of the analyses in this study.

### S1 Detailed Description of Data Acquisition and Curation

#### S1.1 Epidemiological Data

##### COVID-19 case numbers

We obtained the number of confirmed COVID-19 cases from five major sources: (1) the WHO COVID-19 situation reports (WHO 2020), (2) the COVID-19 dashboard published on a Chinese medical website, Ding Xiang Yuan (DXY), that integrates data from local governmental reports (DXY 2020), (3) the National Health Commission of the People's Republic of China (NHCPRC) COVID-19 situation reports (NHCPRC 2020), (4) the European Centre for Disease Prevention and Control (ECDC) COVID-19 situation update (ECDC 2020), and (5) the US Centers for Disease Control and Prevention (USCDC) COVID-19 data tracker (USCDC 2020). Rather than directly collecting data from these sources, we accessed them via two intermediate portals: the `R` (R Core Team 2020) package `nCov2019` (Wu et al. 2020b), and the COVID-19 Data Repository by the Center for Systems Science and Engineering at Johns Hopkins University (Dong et al. 2020).

##### Intervention measures

We focused on two types of intervention measures enacted during the early phase of the COVID-19 pandemic that involved China: targeted-containment measures involving China (*i.e.*, international air-travel bans), and domestic-mitigation measures within China. We compiled information on these containment measures from various news reports, Wikipedia pages (Wikipedia 2020a,b), and peer-reviewed publications (Kraemer et al. 2020; Tian et al. 2020; Hsiang et al. 2020; Lai et al. 2020). For domestic measures within China, we focused on measures that were likely to interrupt travel among regions, including lockdowns at city or province levels, inter-city travel restrictions, and home or neighborhood isolation. See `international_airtravelban_withchina.csv` for a collection of countries or territories that enacted international travel bans with China (and the associated initiation date), and `china_domestic.csv` for a collection of provinces or cities that enacted mitigation measures in China (including the associated implementation period and measure type); these spreadsheets can be found in the GitHub and Dryad repositories.

#### S1.2 Travel Data

##### Global air-travel-volume data

We acquired global air-travel-volume data from FlightAware, containing the number of all commercial passenger flights (subdivided by each aircraft type) per day between December 30, 2019 and March 8, 2020. To estimate the daily number of air-travel passengers, we first gathered information on the number of seats available on each type of aircraft, and then multiplied the number of flights for each type of aircraft by the corresponding number of seats, and then summed over the number of flights for all aircraft each day (shown in Fig. 1B, orange dashed line). The daily number of commercial passenger flights for each type of aircraft is detailed in the report `nflights_daily_byaircraft.csv`, and details on the number of seats for each type of aircraft are provided in `aircraft_nseats.csv` (these spreadsheets can be found in the GitHub and Dryad repositories).

##### China domestic travel-volume data

We used the Baidu Migration dataset (Baidu Inc. 2020; Wang et al. 2014) to assess the volume of domestic travel in China during this early phase of the pandemic and its association with domestic mitigation measures. This dataset is derived from Baidu location-based services (a company whose mapping service holds over 30% market share in China), providing three types of mobility indices that characterize inter-city inflow volume, inter-city outflow volume, and intra-city movement volume. Specifically, we used the aggregated daily domestic mobility index of China, ranging from January 1 to March 8, 2020 (data were unavailable for November and December, 2019). Baidu does not provide information on the relationship between this index and the number of passengers; however, Yuan et al. (2020) estimated that these indices represent approximately 56,137 passengers per unit. Therefore, we used this value to translate the indices into absolute passenger number per day (shown in Fig. 2B, orange dashed line). The daily domestic mobility index of China is contained in `baidu_mobility_index.csv`, which can be found in the GitHub and Dryad repositories.

#### US domestic travel-volume data

We used two publicly available datasets, Apple COVID-19 mobility trends reports (Apple Inc. 2020) and Google COVID-19 community mobility reports (Google LLC 2020) to investigate the domestic-travel volume in the US during the early phase of the pandemic. Apple mobility trends reports are based on direction requests in the Apple Maps app, and provide three types of daily mobility indices: walking, driving, and transit. Each daily mobility index is the relative number of direction requests made on that day compared to the corresponding baseline volume on January 13, 2020 (these indices are only made available by Apple from this date onward to help study mobility trends during the COVID-19 pandemic). Google community mobility data are collected from smartphones with the Google location-based services turned on. These mobility data recognize six types of locations: (1) transit; (2) retail and recreation; (3) groceries and pharmacy; (4) parks; (5) workplaces, and; (6) residential. These mobility indices are relative values, where the mobility index for a given day is computed as the number of visits on that day divided by the median number of visits on that day measured from January 3 to February 6, 2020 (these indices are only made available by Google from February 6 onward onward to help study mobility trends during the COVID-19 pandemic). The daily domestic mobility indices of the US from Apple and from Google are contained in `apple_mobility_indices.csv` and `google_mobility_indices.csv`, respectively (these spreadsheets can be found in the GitHub and Dryad repositories).

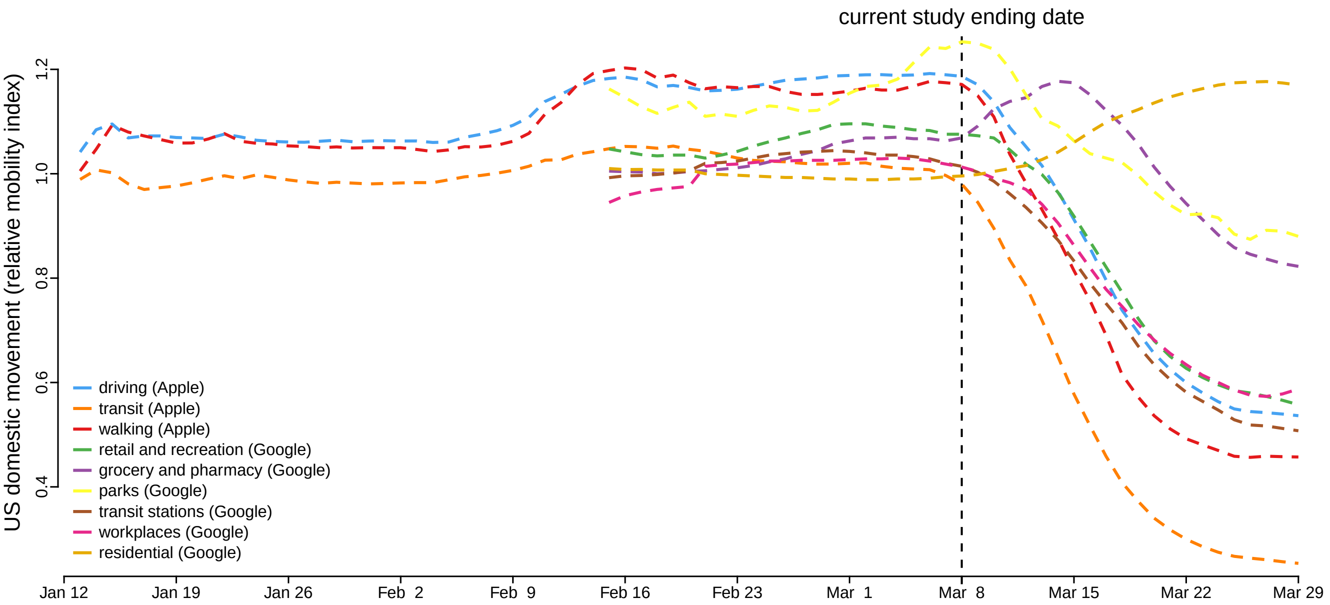


Figure S2: **US domestic mobility indices.** (7-day smoothed average) US domestic mobility indices acquired from different sources consistently indicate that domestic measures drastically reduced population movement within the US, but they were not enacted (or had no visible impact) until the week of March 9, 2020.

#### S1.3 Definition of Time Intervals

We partitioned the early phase of the COVID-19 pandemic into four time intervals of similar duration: (1) interval 1 from late 2019 (origin time) to January 12, 2020; (2) interval 2 from January 13, 2020 to February 2, 2020; (3) interval 3 from February 3, 2020 to February 16, 2020, and; (4) interval 4 from February 17, 2020 to March 8, 2020.

The boundary between the first and second intervals coincides with the start of the Spring Festival travel season in China. The boundary between the second and third intervals (February 2) coincides with the initiation of international air-travel bans with China (imposed by 34 countries by this date) and the cancellation or significant reduction of international air services involving China (by over 130 airlines; International Civil Aviation Organization 2020; Wikipedia 2020b), following the declaration of Public Health Emergency of International Concern (PHEIC) by the World Health Organization (WHO) on January 31, 2020. This boundary also falls close to the onset of widespread mitigation measures in China to restrict domestic travel: these measures began with the lockdown of Wuhan on January 23, 2020 (extended to the entire Hubei province in the following days), followed by the declaration of level-1 emergency in all mainland provinces between January 24–29, 2020, the extension of the Spring Festival national holiday (effectively school and workplace closure) announced on January 27, and the enactment of stringent home- or neighborhood-isolation orders in various cities outside Hubei beginning February 2, 2020. The boundary between the third and fourth intervals coincides with the lifting of travel restrictions in China (except in Hubei, where the travel restrictions were not lifted until late March, 2020).

#### S1.4 SARS-CoV-2 Genomic Sequence Data

We curated two SARS-CoV-2 genomic sequence datasets for our study, one with 1271 sequences (the "reduced dataset") and the other with 2598 sequences (the "entire dataset").

##### Assembling the reduced dataset

The reduced dataset consists of all available SARS-CoV-2 genomic sequences available as of April 19, 2020 from GISAID (`https://www.gisaid.org/`; Shu and McCauley 2017). As our focus is on the crucial early phase of the COVID-19 pandemic, we excluded sequences that were collected after March 8, 2020, leaving 2003 sequences in the dataset. We first filtered the dataset by excluding sequences that fit any of the following conditions: (1) fewer than 29000 sites (not counting missing or gap sites); (2) lacking associated metadata (*e.g.*, sampling time or location); (3) lacking the precise sampling date or geographic location (state/province for sequences from China, Canada, or U.S.A. and country for the others); (4) sampled from a non-human host; (5) multiple sequences from the same individual (in which case we randomly selected one of sequence and discarded the others), or; (6) duplicates of other sequences in the dataset [for this purpose, we used the "exclude list" used by `Nextstrain` (`https://github.com/nextstrain/ncov/blob/master/defaults/exclude.txt`) as a reference]. Application of these filters resulted in a genomic dataset consisting of 1620 sequences.

We then inferred an alignment of these nucleotide sequences using `MUSCLE` version 3.8 (Edgar 2004). We performed a second round of filtration of the resulting alignment. First, we excluded sequences that appeared to be anomalously divergent; this was achieved by comparing each sequence to the reference genome while assuming that the rate of mutation accumulation should not exceed 10 mutations per genome per month. We also excluded sequences with many ambiguous sites (*i.e.*, sites for which the nucleotide could not be unambiguously identified); specifically, we discarded sequences with more than 15 ambiguous sites, and sequences with at least 10 ambiguous sites and fewer than 10 sites differing from the reference genome. Next, we excluded sequences with nonsense mutations; to this end, we translated the nucleotide alignment into an amino-acid alignment using the `seqinr` package (Charif and Lobry 2007) in `R` (R Core Team 2020) to identify sequences with premature stop codons. We assumed that the rate of amino-acid substitution accumulation should not exceed 4 substitutions per genome per month; we therefore discarded sequences with more than 6 ambiguous amino-acid sites, and sequences with at least 3 ambiguous amino-acid sites but fewer than 3 sites differing from the reference genome. After filtering, our reduced dataset included 1271 sequences.

#### Assembling the entire dataset

We also compiled a more comprehensive dataset by curating all sequences available from GISAID as of September 22, 2020. Specifically, we downloaded an alignment from GISAID, which was inferred using `MAFFT` (Katoh and Standley 2013). After excluding sequences that were collected after March 8, 2020, the alignment included 4012 sequences. We then performed two filtration steps identical to those we performed on the reduced dataset, culminating in an alignment ("entire dataset") with 2598 sequences.

The entire dataset is more comprehensive than the reduced dataset: it contains more than twice the number of sequences (Fig. S3), and is also more evenly sampled, as the sequence-to-case ratios of many undersampled geographic areas are significantly higher, especially for the third and fourth intervals of our study (Fig. S4). Moreover, the entire dataset contains SARS-CoV-2 genomic sequences that are likely to represent the vast majority of such data that will ever be available; the deposition rate of sequences collected from the early phase of the pandemic drastically decreased in September, 2020 (Fig. S3).

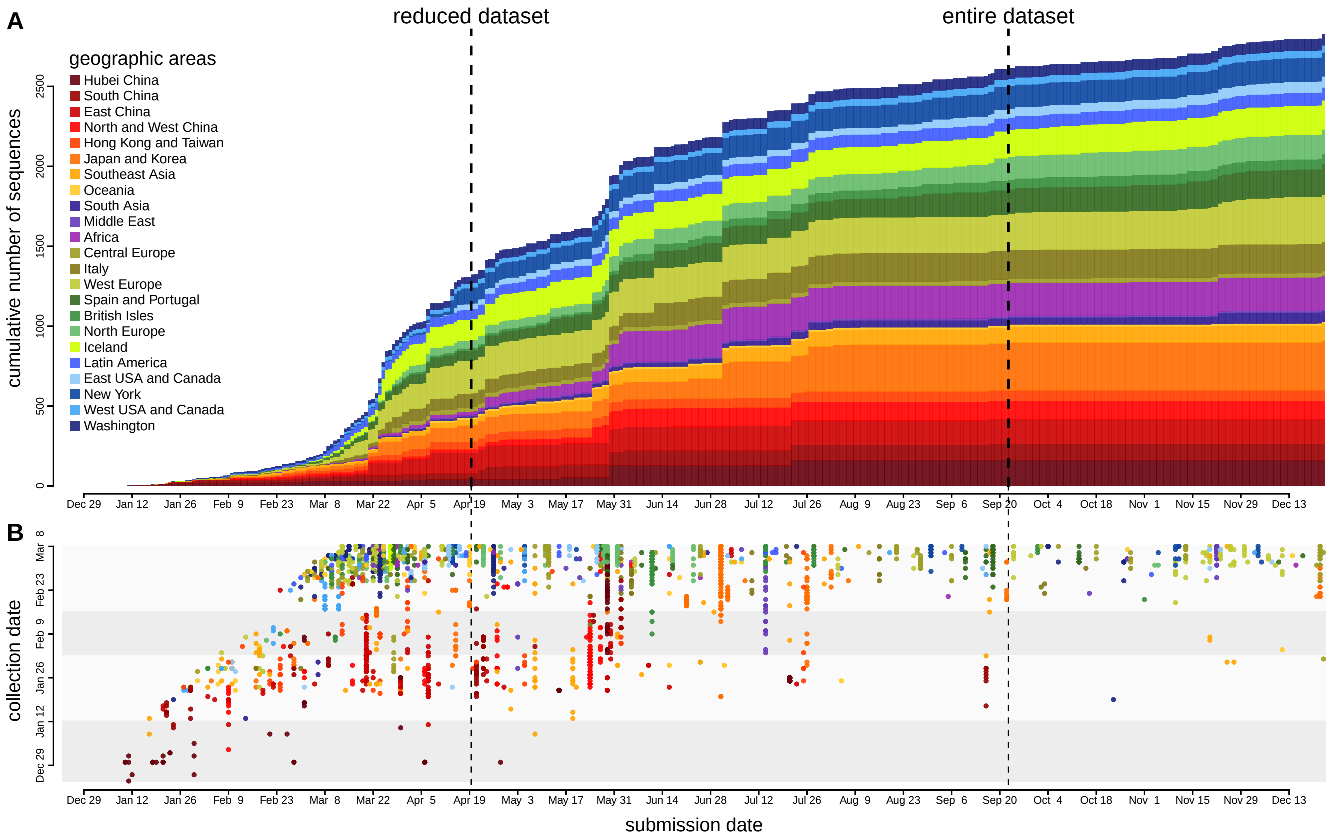


Figure S3: **Submission and collection dates for SARS-CoV-2 genomic sequences.** **(A)** Cumulative number of SARS-CoV-2 sequences submitted to GISAID that were collected during the early phase of the COVID-19 pandemic. The color of each segment in the stacked bar plot indicates the number of sequences submitted from the corresponding geographic area on that day. **(B)** Submission and collection dates for each SARS-CoV-2 sequence included in our study. The deposition rate of sequences collected prior to March 8 drastically decreased in September, 2020.

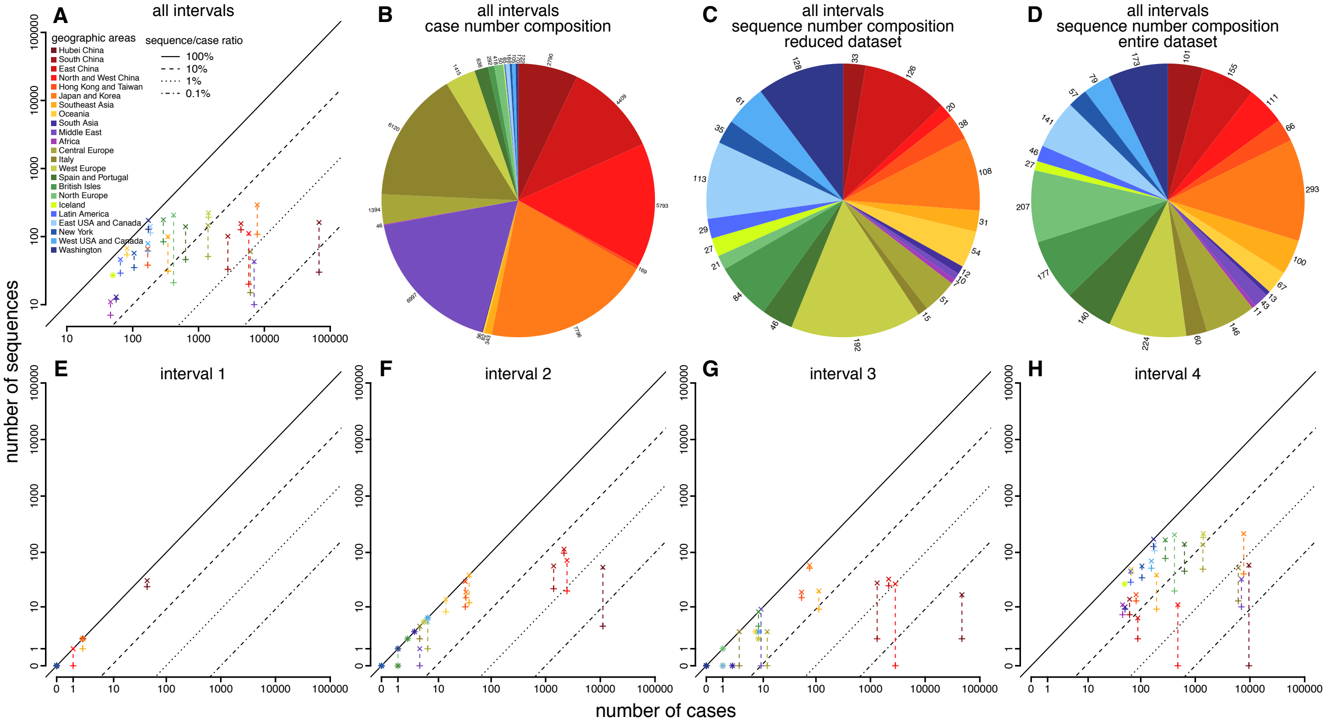


Figure S4: **Number of SARS-CoV-2 genome sequences versus number of confirmed COVID-19 cases. (A)** Number of sequences versus number of confirmed cases across the early phase of the pandemic. + and x indicate the reduced and entire datasets, respectively; they are connected by dashed line for each geographic area to show the increase of the number of sequences (and thus the sequence/case ratio) in the entire dataset. **(B)** Geographic distribution of confirmed case numbers (excluding Hubei). **(C)** Geographic distribution of sequence number (excluding Hubei; reduced dataset). **(D)** Geographic distribution of sequence number (excluding Hubei; entire dataset). **(E–H)** Number of sequences versus number of confirmed cases for each interval of the early phase, respectively.

#### Trimming and partitioning the curated alignment

For each curated dataset, we trimmed the 5′UTR and 3′UTR as well as the other non-coding regions, retaining only coding regions in the alignment. Table S1 lists the coding regions and their corresponding coordinates in the reference genome (Wuhan-Hu-1, Wu et al. 2020a). After removing the stop codon for each coding region, both the reduced and entire alignments for the complete coding region included 29,232 nucleotide sites.

Table S1: **Genomic coordinates of the SARS-CoV-2 coding regions.**

| Region | Starting coordinate | Ending coordinate |
|---|---|---|
| ORF1ab* | 266 | 21555 |
| S | 21563 | 25384 |
| ORF3a | 25393 | 26220 |
| E | 26245 | 26472 |
| M | 26523 | 27191 |
| ORF6 | 27202 | 27387 |
| ORF7a | 27394 | 27759 |
| ORF7b | 27756 | 27887 |
| ORF8 | 27894 | 28259 |
| N | 28274 | 29533 |
| ORF10 | 29558 | 29674 |

*During translation, ORF1ab experiences a −1 ribosomal frameshift at site 13468, so the range is (266–13468, 13468–21555).

### S2 Detailed Description of Phylodynamic Analyses

#### S2.1 Estimating a Dated Phylogeny for the Reduced SARS-CoV-2 Dataset

##### Overview

In this section, we describe the analyses that we performed to infer a dated phylogeny for the reduced sample of COVID-19 viruses. The phylogeny inferred from these analyses will be used in our subsequent evaluation of candidate biogeographic models in Section S2.2.

##### Model specification

We inferred a dated phylogeny by performing Bayesian analyses of the reduced SARS-CoV-2 sequence dataset under a relaxed-clock model, which includes three main components: (1) a substitution model; (2) a branch-rate prior model; and (3) a branching-process prior model. Below, we describe each of these model components and the corresponding priors for the parameters of those models (note that we used an empirical Bayesian approach to specify non-default priors for several parameters; *i.e.*, where the results of preliminary analyses and/or published results were used to specify the parameters of priors. Details of the priors are described in Table S2.

###### Substitution model

The substitution model collectively describes the process of molecular evolution of the SARS-CoV-2 genomes over the branches of the phylogeny. The process of molecular evolution is apt to vary among regions of these viral genomes. For example, the ORF1ab gene of SARS-CoV-2 encodes nonstructural proteins and is therefore likely to have been subjected to strong purifying selection (Li et al. 2020b), whereas other genes, such as the spike (S) gene, encode structural proteins that determine antigenicity and other immune properties of SARS-CoV-2, and are therefore likely to have been subjected to strong positive selection (Korber et al. 2020; Plante et al. 2020; Hou et al. 2020; Volz et al. 2021). Accordingly, we specified a partitioned substitution model to accommodate possible variation in the evolutionary process across genomic regions. Specifically, we partitioned the SARS-CoV-2 genomes into six data subsets, with three subsets for the ORF1ab gene region (one for each codon position), and three subsets for the remaining ten combined gene regions (one for each codon position). (For a complete list of the gene regions and their corresponding coordinates in the reference genome, see Table S1).

For each of these data subsets, we specified an independent TN93 substitution model (Tamura and Nei 1993), with transition-transversion rate-ratio parameters $\kappa_1$ and $\kappa_2$ (the instantaneous rates of A to G and C to T substitutions, respectively, relative to the transversion rate) and $\pi$ (the stationary frequency

Table S2: **Priors used to estimate a dated phylogeny of the sampled SARS-CoV-2 sequences.**

| Parameter | Description | Prior |
|---|---|---|
| $\kappa_1$ | Ratio of the A $\to$ G rate to the transversion rate | Lognormal$(\mu = 1.0, \sigma = 0.8)^*$ |
| $\kappa_2$ | Ratio of the C $\to$ T rate to the transversion rate | Lognormal$(\mu = 1.0, \sigma = 0.8)$ |
| $\boldsymbol{\pi}$ | Nucleotide stationary frequencies | Dir$(1, 1, 1, 1)$ |
| $\boldsymbol{m}$ | Partition-specific rate multipliers | Dir$(1, 1, 1, 1, 1, 1)$ |
| $\alpha$ | Shape and scale parameter of the $\Gamma_4$ distribution | Lognormal$(\mu = -2.1, \sigma = 0.5874)$ |
| $\mathbb{E}[\boldsymbol{r}]$ | Mean of the UCLN | Lognormal$(\mu = -12.7, \sigma = 0.5874)$ |
| $SD(\boldsymbol{r})$ | Standard deviation of the UCLN | Exp$(\lambda = 1/(2.0\text{e}{-6}))$ |
| $N_T$ | Effective number of infected individuals at sampling time, $T$ | Lognormal$(\mu = 7.5, \sigma = 1.0)$ |
| $r$ | Exponential growth rate of the coalescent model | Laplace$(0.07, 0.01)$ |

*$\mu$ and $\sigma$ in this table are the mean and standard deviation of the normal distribution.

of each nucleotide). For each transition-transversion rate-ratio parameter, we specified lognormal priors with a prior mean of 3.74 and a 95% prior interval of $[0.56, 13.04]$.

To accommodate possible variation in the overall rate of substitution *between* gene regions, we specified independent rate multipliers for each of the six data subsets. To accommodate variation in substitution rates across sites *within* each data subset, we specified a discrete-gamma model (Yang 1994). Our preliminary analyses specified an independent discrete-gamma model for each of the six data subsets, which revealed a similar degree of among-site rate variation within each data subset (*i.e.*, with similar posterior estimates for the six $\alpha$-shape parameters). Accordingly, to decrease model complexity (and reduce MCMC issues), we specified a shared, discrete-gamma model for the entire alignment. We specified a lognormal hyperprior on the $\alpha$-shape parameter, with a prior mean of 0.15 and a 95% prior interval spanning one order of magnitude around the mean, $[0.039, 0.39]$. This prior reflects our expectation of a high degree of substitution-rate variation across sites, motivated by our observation that most sites in our SARS-CoV-2 alignment are invariant, while a small number of sites appear to be highly variable.

**Branch-Rate model**

The branch-rate model describes how the overall substitution rate varies across branches of the tree. Our composite relaxed-clock model specifies the uncorrelated lognormal (UCLN) branch-rate prior model (Drummond et al. 2006; Li and Drummond 2012; Rannala and Yang 2007), which accommodates variation in the overall substitution rate across branches by drawing i.i.d. rate multipliers for each branch from a shared underlying lognormal distribution, where the parameters of this distribution (mean and standard deviation) are estimated from the data. For the mean of the UCLN, we specified a lognormal hyperprior with an expectation of $3.63\text{e}{-6}$ substitutions/site/day and 95% prior interval of $[0.96\text{e}{-6}, 9.64\text{e}{-6}]$, motivated by published substitution-rate estimates of approximately 30 substitutions/genome/year (*cf.* Duchene et al. 2020). We specified an exponential hyperprior on the standard deviation of the UCLN such that the branch-specific substitution rates are expected to vary over approximately one order of magnitude.

**Branching-Process model**

The branching-process model describes the prior distribution of tree topologies and divergence times. We used a coalescent model with exponential population growth as our branching-process model. This model assumes that the viral population size grows as a deterministic exponential function (Beaumont 1999; Drummond et al. 2002), which is motivated by the fact that our SARS-CoV-2 dataset was sampled from the early, explosive stage of the COVID-19 pandemic. This model is completely described by two free parameters: $N_T$, the effective number of infected individuals in the population at the sampling time, $T$ (*i.e.*, the last sampling date in our dataset, March 8, 2020), and $r$, the exponential growth rate. We specified empirically informed and biologically realistic priors on these parameters.

*Prior on the exponential growth rate, r*

We specified a prior on $r$ using external information about the $R_0$, the basic reproductive number, and $\tau$, the duration of the infectious period; these quantities are related through the equation $r = (R_0 - 1)/\tau$. We specified a Laplace prior on $r$, with location and scale parameter values specified according to published estimates of $R_0$ and $\tau$ for COVID-19 (Chinazzi et al. 2020; Li et al. 2020a; Hao et al. 2020; Vaughan et al. 2020; Nadeau et al. 2021; Wölfel et al. 2020; van Kampen et al. 2021; Byrne et al. 2020). This prior on $r$ directly translates to an expected population doubling time, $t = \ln(2)/r$, of 10.6 days (with 95% prior interval ranging from 6.9 to 17.2 days).

*Prior on the effective population size, $N_T$*

Given a population doubling time of $t$, the expected number of individuals at time $T$ is $N_T = N_0 2^{(T-T_0)/t}$, where $N_0$ is the number of individuals at the beginning of the process, and $T_0$ is the origin time of the process. (This equation follows from the fact that there are $(T - T_0)/t$ doubling cycles in a period of duration $T - T_0$.) We therefore specified a prior on $N_T$ informed by our previously determined prior on $t$, as well as several realistic values for $N_0$ (1 or 2) and $T_0$ (some time in late November, 2019). Based on these values, we chose a lognormal prior on $N_T$ such that the mean was 2981 and 95% prior interval spanned $[254, 12840]$. Given that there were at least 3940 reported cases on March 8, 2020 alone, it may seem unreasonable to specify a prior such that the expected number of individuals is as low as 2981. However, we note that $N_T$ represents the effective number of infected individuals in the population, which is typically substantially smaller than the total number of infected individuals in the population. Additionally, we note that: (1) the posterior-mean estimate of $N_T$ under this prior is $\approx 801$, indicating that, if anything, this prior mean is too high, and; (2) sensitivity analyses suggested that posterior estimates of $N_T$ were not very sensitive to this prior (results not shown).

##### Parameter estimation

We performed six independent MCMC simulations to approximate the joint posterior distribution of the relaxed-clock model parameters using `BEAST` version 1.10.5 (Suchard et al. 2018) with the `BEAGLE` library (compiled from the 'hmc-clock' branch, commit 'dd36bf5': `https://github.com/beagle-dev/beagle-lib/tree/dd36bf5b8d88348c77a93eeeef0917d90df71a4f`; Ayres et al. 2019) enabled to accelerate computation. We ran each replicate MCMC simulation for 50 million generations, sampling continuous parameters every 1000 generations and trees every 10,000 generations. Details of these analyses (*e.g.*, proposal weights) are available in the XML scripts included in our GitHub and Dryad repositories. After discarding the first 20% as the burn-in from each replicate simulation, we combined the remaining posterior samples of trees from all the replicates and then down-sampled every 50,000 generations using `LogCombiner` version 1.10.5. Following initial inspection of the log files using Tracer (Rambaut et al. 2018) version 1.7.1, we further evaluated MCMC performance using the `coda` package (Plummer et al. 2006) in `R` (R Core Team 2020). We assessed convergence of replicate MCMC simulations by calculating the ESS for each continuous parameter for the combined posterior samples; ensuring that values for the substitution-model parameters were all $\gg 10000$ and those for the branch-rate and branching-process models were all $\gg 200$. We then used `TreeAnnotator` version 1.10.5 to generate a summary phylogeny from the combined posterior sample of trees—as a maximum clade credibility (MCC) tree—where the age of each internal node is computed by marginalizing over the age of that node across all samples. Note that as the age of each node is summarized independently across the posterior distribution of trees, it is possible for the MCC summary tree to have negative branch lengths (*i.e.*, where a descendant node is older than its ancestor). To avoid potential issues caused by this phenomenon in downstream analyses, we assigned a small positive value (0.001 days) as the duration of these "time-traveling" branches.

#### S2.2 Evaluating Candidate Biogeographic Models

##### Overview

In this section, we describe our analyses to explore candidate biogeographic models that describe the geographic progression of the SARS-CoV-2 virus during the early phase of the COVID-19 pandemic. We begin by briefly describing the general features of these biogeographic models, and then define the space of candidate models that we will evaluate. We then describe the analyses that we performed to both assess the *relative fit* of these candidate biogeographic models to our reduced SARS-CoV-2 dataset (using Bayes factors), and to assess the *absolute fit* of each candidate biogeographic model to our dataset (using posterior-predictive simulation). In evaluating candidate biogeographic models, we condition on the MCC summary phylogeny for the reduced SARS-CoV-2 sequence dataset described in Section S2.1.

##### Biogeographic models

###### Generic description of discrete biogeographic models

Biogeographic models describe the history of geographic dispersal over the tree, $\Psi$, as a continuous-time Markov chain (CTMC). For a biogeographic history with $k$ discrete areas, this stochastic process is fully specified by a $k \times k$ instantaneous-rate matrix, $\mathbf{Q}$, where an element of the matrix, $q_{ij}$, is the instantaneous rate of change between state $i$ and state $j$ (*i.e.*, the instantaneous rate of dispersal from area $i$ to area $j$). Each element, $q_{ij}$, of the instantaneous-rate matrix, $\mathbf{Q}$, is specified as:

$$q_{ij} = r_{ij}\delta_{ij},$$

where $r_{ij}$ is the relative rate of dispersal between areas $i$ and $j$, and $\delta_{ij}$ is an indicator variable that takes one of two states (0 or 1). When $\delta_{ij} = 1$, the instantaneous dispersal rate for the corresponding element, $q_{ij}$, is simply $q_{ij} = r_{ij}$. Conversely, when $\delta_{ij} = 0$, the instantaneous dispersal rate for the corresponding element, $q_{ij}$, is zero, effectively removing that parameter from the biogeographic model. Each unique vector of $\delta_{ij}$—*i.e.*, $\delta$, a string of zeros and ones for each of the possible pairwise dispersal routes between the $k$ geographic areas—corresponds to a unique biogeographic model. The total number of dispersal routes for a given biogeographic model is denoted $\Delta$. By convention, we rescale the $\mathbf{Q}$ matrix such that the expected number of dispersal events in one time unit is equal to the parameter $\mu$ (Yang 2014).

###### Space of candidate biogeographic models

*Symmetry of the instantaneous-rate matrix*

Alternative biogeographic models may be specified based on the symmetry of the instantaneous-rate matrix. The biogeographic model described by Lemey *et al.* (Lemey et al. 2009) assumes the instantaneous-rate matrix, $\mathbf{Q}$, is symmetric, where $q_{ij} = q_{ji}$ (*i.e.*, $r_{ij} = r_{ji}$ and $\delta_{ij} = \delta_{ji}$); namely, the instantaneous rate of dispersal from area $i$ to area $j$ is assumed to be equal to the dispersal rate from area $j$ to area $i$. A subsequent extension (Edwards et al. 2011) allows the $\mathbf{Q}$ matrix to be asymmetric, *i.e.*, $q_{ij}$ and $q_{ji}$ are not constrained to be equal, allowing the rate of dispersal from area $i$ to area $j$ to be different from the rate of dispersal from area $j$ to area $i$.

*Prior on the number of dispersal routes*

Recall that the total number of dispersal routes for a given biogeographic model is $\Delta$. The default prior in `BEAST`, proposed by in Lemey *et al.* (Lemey et al. 2009), reflects a preference for biogeographic models with the minimal number of dispersal routes (Table S3). Specifically, the default prior on $\Delta$ for the symmetric model is an *offset* Poisson prior that assigns zero probability to all biogeographic models with fewer than $k - 1$ dispersal routes. This prior reflects the constraint that a dataset with $k$ geographic areas cannot be realized under a CTMC with fewer than $k - 1$ non-zero $q_{ij}$ values (*i.e.*, dispersal routes). The prior on the number of dispersal routes *greater than* $k - 1$ is described by a Poisson prior with rate parameter, $\lambda$. By default, $\lambda = \ln(2)$, which places 50% of the prior probability on biogeographic models with the minimum number of dispersal routes, *i.e.*, where $\Delta = (k-1)$. For the asymmetric model (Edwards et al. 2011), the number of dispersal routes is assumed to be drawn from a Poisson prior with rate $\lambda$. In this case, $\lambda$ is specified such that the expected number of dispersal routes is $k-1$ (note that this prior does not enforce a minimum number of dispersal routes).

We also consider an alternative and more diffuse prior on the total number of dispersal routes, $\Delta$ (Table S3). This alternative prior is specified by setting the expected number of dispersal routes to be about half the maximum number; this results in a relatively flat prior probability that any given dispersal route exists for all values of $k$. Specifically, for the symmetric model, we specify an offset (*i.e.*, by $k-1$) Poisson prior on $\Delta$ with $\lambda$ specified so that the expected number of dispersal routes is about half of the maximum number, $\binom{k}{2}$, for a dataset with $k$ areas. For the asymmetric model, we specify a Poisson prior distribution on $\Delta$ with $\lambda = \binom{k}{2}$, which represents a prior belief that half of all possible dispersal routes are included in the biogeographic model.

*Prior on the average dispersal rate*

Recall that the rate matrix, $\mathbf{Q}$, is rescaled so that the average rate of dispersal between all areas is $\mu$. For a tree of length $T$ (*i.e.*, the sum of the durations of all branches in the tree), the expected number of dispersal events is $\mu \times T$. Therefore, the prior on $\mu$ represents our prior belief about the number of dispersal events over the tree. By default, the prior on $\mu$ implemented in `BEAST` is a gamma prior with shape parameter $\alpha = 0.5$ and rate parameter $\beta = T$. The gamma distribution has a mean of $\alpha/\beta$; therefore this prior expresses the belief that the average dispersal rate is $0.5/T$. Because the expected number of dispersal events is $\mu \times T$, the expected number of dispersal events under this prior is 0.5, independent of the duration of the entire biogeographic history (*i.e.*, the tree length, $T$), or the number of areas, $k$, in which the pathogen occurs. Accordingly, the default prior on $\mu$ in `BEAST` reflects a strong prior expectation that the biogeographic history under study involved a low average dispersal rate and a very small number of dispersal events.

We consider an alternative and more permissive prior on the average dispersal rate, $\mu$ (Table S3). This alternative specifies an exponential prior on $\mu$ with rate parameter $\theta$, and a mean of $1/\theta$. Rather than assuming a fixed value for the mean of the exponential prior, we treat it as a random variable to be estimated from the data. Specifically, we specify a gamma hyperprior on $1/\theta$; this gamma hyperprior has shape parameter $\alpha = 0.5$ and rate parameter $\beta = 0.5$ (enforcing the shape and rate parameters to be equal ensures that the resulting prior on $\mu$ is proper). The resulting prior—known as the $K$-distribution (Jakeman and Pusey 1978)—is more diffuse than the default prior on $\mu$, as is the resulting prior distribution on the number of dispersal events.

Table S3: **Priors used in evaluating candidate biogeographic models.**

| Parameter | Model | Default | Alternative |
|---|---|---|---|
| Number of dispersal routes, $\Delta$ | Symmetric | $(\Delta - k + 1) \sim \text{Pois}(\ln 2)$ | $(\Delta - k + 1) \sim \text{Pois}(\lceil \frac{k^2-5k+4}{4} \rceil)$ |
| | Asymmetric | $\Delta \sim \text{Pois}(k-1)$ | $\Delta \sim \text{Pois}(\frac{k(k-1)}{2})$ |
| Average dispersal rate, $\mu$ | — | $\mu \sim \Gamma(0.5, T)$ (*i.e.*, CTMC-rate reference) | $\mu \sim \text{Exp}(1/\lambda)$ $\lambda \sim \Gamma(0.5, 0.5)$ |
| Relative dispersal rate, $r_{ij}$ | | $\Gamma(1,1)$ | $\Gamma(1,1)$ |

*Constant vs piecewise-constant dispersal rate and rate matrix*

We can specify alternative biogeographic models based on the assumed constancy of the dispersal process. For example, the simplest possible model assumes that the average dispersal rate and the dispersal dynamics (*i.e.*, the instantaneous-rate matrix) remain constant over the entire biogeographic history. More complex, piecewise-constant biogeographic models allow the average dispersal rate and/or the dispersal dynamics (*i.e.*, the instantaneous-rate matrix) to differ between two or more pre-specified intervals of the geographic history. Bielejec and colleagues (Bielejec et al. 2014) developed a piecewise-constant model (Lemey et al. 2009) that allows the instantaneous-rate matrices to vary among two or more pre-specified time intervals (but assumes that the average dispersal rate is constant across intervals). Subsequently, Membrebe and colleagues (Membrebe et al. 2019) developed a piecewise-constant model that allows the average dispersal rate to vary among two or more pre-specified time intervals (but assumes that dispersal dynamics remain constant across intervals). Here, we evaluate biogeographic models that allow both the average dispersal rate and the dispersal dynamics (instantaneous-rate matrix) to vary in a piecewise-constant manner.

Specifically, this piecewise-constant model partitions the entire history into $m$ time intervals separated by $m+1$ breakpoints dated at $\boldsymbol{T} = \{T_0, T_1, \ldots, T_{l-1}, T_l, \ldots, T_{m-1}, T_m\}$, where $T_0 = -\infty$, $T_{l-1} < T_l$, and $T_m = 0$; *i.e.*, the first interval spans a period reaching deep into the past until $T_1$, and the most recent interval spans $T_{m-1}$ to the present (the end of the sampling period). This model is piecewise constant in that the instantaneous-rate matrix, $\mathbf{Q}$, and the average dispersal rate, $\mu$, are constant within each interval, but are free to vary independently between the pre-specified intervals. Therefore, the dispersal process is fully specified by two vectors: $\vec{\mathbf{Q}} = \{\mathbf{Q}_1, \ldots, \mathbf{Q}_m\}$ and $\boldsymbol{\mu} = \{\mu_1, \ldots, \mu_m\}$. See Bielejec et al. 2014; Membrebe et al. 2019; Landis 2017 for further details regarding the model specification.

#### Evaluating candidate biogeographic models

We explored a pool of candidate biogeographic models corresponding to 12[1] possible combinations of: (1) symmetric and asymmetric $\mathbf{Q}$ matrices; (2) default and alternative priors on the total number of dispersal routes; (3) default and alternative priors on the average dispersal rate, and; (4) models where the $\mathbf{Q}$ matrices and average dispersal rates are piecewise constant over 1, 2, or 4 pre-specified intervals. The 2-interval models set a boundary on February 2, 2020 (the onset of international air-travel ban with China), the 4-interval models set boundaries on January 12, 2020 (the start of the Chinese Spring Festival), February 2, and February 16, 2020 (the cessation of domestic-travel restrictions in China); see Section S1.3 for details. We assessed both the *relative fit* of these candidate biogeographic models to our reduced SARS-CoV-2 dataset (by computing Bayes factors to compare competing models), and also assessed the *absolute fit* of each candidate biogeographic model to our SARS-CoV-2 dataset (using posterior-predictive simulation).

##### Assessing relative fit of candidate biogeographic models using Bayes factors

We evaluated the relative fit of each candidate biogeographic model to our SARS-CoV-2 dataset using Bayes factors. This Bayesian model-comparison approach requires that we first estimate the marginal likelihood for each candidate biogeographic model, and then compute the Bayes factor for each pair of competing models as twice the difference in their log marginal likelihoods (Kass and Raftery 1995). We estimated marginal likelihoods for each candidate biogeographic model using both thermodynamic-integration (Lartillot and Philippe 2006) and stepping-stone (Xie et al. 2011; Baele et al. 2012) estimators. These marginal-likelihood estimators tend to be unstable when inferring the phylogeny and biogeographic history jointly, owing to the diffuse (hyper)priors on node-age and branch-rate model parameters, as well as the vast tree space (see Baele et al. 2015). Accordingly, we estimated marginal likelihoods for our candidate biogeographic models by conditioning on the summary phylogeny (the MCC tree) that we inferred using sequence data alone (see Section S2.1).

For each candidate biogeographic model, we ran four replicate power-posterior MCMC simulations implemented in `BEAST` (Suchard et al. 2018) with the `BEAGLE` library (version 3.2.0; Ayres et al. 2019). Specifically, the analyses under the time-constant biogeographic models were performed using `BEAST`

[1]Note that we excluded the remaining 12 possible candidate biogeographic models from rigorous evaluation because they involved combinations of individual model components that generally provided a poor fit to our SARS-CoV-2 datasets.

version 1.10.5, whereas those under the piece-wise constant biogeographic models were performed using our modified version of BEAST (see Section S3 for details). For each replicate power-posterior MCMC simulation, we used 36–48 powers placed at evenly-spaced quantiles of a $\text{Beta}(0.3, 1.0)$ distribution. For each power, we discarded the initial 65000–100000 generations as burn-in and then sampled every 100 generations in the remaining 210000–250000 generations. We assessed the reliability of our marginal-likelihood estimates by comparing values from the four replicate power-posterior simulations. Details of these analyses (*e.g.*, proposal weights) are available in the XML scripts included in our GitHub and Dryad repositories.

**Assessing absolute fit of candidate biogeographic models using posterior-predictive simulation**

We assessed the absolute fit of each candidate biogeographic model to our SARS-CoV-2 biogeographic dataset using posterior-predictive simulation (Gelman et al. 1996). In principle, posterior-predictive simulation is based on the following premise: if a given model provides an adequate description of the true process that gave rise to our observed data, then datasets simulated under that model should resemble our observed dataset. In practice, posterior-predictive simulation involves the following procedure: (1) we first estimate the joint posterior probability distribution of parameters for the candidate model from the observed biogeographic dataset; (2) then we randomly draw a vector of sampled parameter values from the inferred joint posterior distribution, where the sample consists of a fully specified biogeographic model; (3) for each sample, we simulate a new biogeographic dataset on the sampled tree given the sampled parameters of the biogeographic model; (4) finally, we compute a summary statistic that measures the discrepancy between the observed dataset and the simulated dataset. We repeat steps 2–4 many times to generate a distribution of the summary statistic that is predicted from the posterior of the candidate biogeographic model (*i.e.*, a *posterior-predictive distribution*). If the candidate biogeographic model provides an adequate description of the process that gave rise to our observed data, the resulting posterior-predictive distribution should contain zero with high probability. We assessed model adequacy using two summary statistics (the parsimony and tip-wise multinomial statistics), which we describe in Section S3.3.

*Estimating the joint posterior probability distribution for each candidate biogeographic model*

For each of the candidate biogeographic models, we first inferred the joint posterior distribution from the observed biogeographic data (*i.e.*, the geographic location of each of the sequences in our reduced SARS-CoV-2 dataset) by performing two to four independent MCMC simulations using BEAST (Suchard et al. 2018) with the BEAGLE library (version 3.2.0; Ayres et al. 2019). Specifically, the analyses under the time-constant biogeographic models were performed using BEAST version 1.10.5, whereas those under the piece-wise constant biogeographic models were performed using our modified version of BEAST (see Section S3 for details). For each replicate MCMC simulation, we ran 10 million generations, sampling every 2000 generations. We discarded the initial 10% of samples (as burn-in) from each replicate MCMC simulation, and then combined the remaining posterior samples from all replicate simulations using LogCombiner version 1.10.5. We then assessed MCMC performance for the resulting composite posterior sample by inspecting the log files using Tracer (Rambaut et al. 2018) version 1.7.1, and using the coda package (Plummer et al. 2006) in R (R Core Team 2020). Specifically, we ensured that the computed ESS values for all continuous parameters were $\gg 100$. Details of these analyses are available in the XML scripts included in our GitHub and Dryad repositories.

*Posterior-predictive simulations*

For each candidate biogeographic model, we simulated $m = 2500$ predictive datasets by repeatedly sampling at random from the corresponding joint posterior probability distribution. We then generated posterior-predictive distributions from each set of $m$ predictive datasets under 10 separate test statistics, *i.e.*, for all combinations of the two (parsimony and tip-wise multinomial) summary statistics, and each of five time intervals (*i.e.*, the entire history and each of the four intervals of the piecewise-constant biogeographic model). For each posterior-predictive distribution, we computed the posterior-predictive p-value (see Section S3.3 for details).

#### Results

Bayes-factor comparisons of all candidate models provide decisive support (*i.e.*, $2 \ln \text{BF} \gg 10$; Table S4) for the 4-interval piecewise-constant asymmetric biogeographic model with alternative priors for both the average dispersal rate and the total number of dispersal routes. The preference for this model is corroborated by the results of our posterior-predictive simulations: the model selected by the Bayes-factor comparisons was inferred to provide an adequate absolute fit to our reduced SARS-CoV-2 dataset for every summary statistic, whereas the other 11 candidate models were inferred to be inadequate by one or more of the summary statistics (Figure S5). Accordingly, we use the 4-interval piecewise-constant asymmetric alternative-prior biogeographic model for our joint phylodynamic analyses of the entire SARS-CoV-2 dataset described below (see Section S2.3).

##### The relative fit of competing biogeographic models to the SARS-CoV-2 dataset

Table S4: **Marginal-likelihood estimates of the candidate biogeographic models conditioning on the MCC phylogeny for the reduced SARS-CoV-2 dataset.** Columns 3–6 list marginal likelihoods inferred from four replicate power-posterior MCMC simulations, column 7 lists the composite marginal-likelihood estimates (computed by combining the replicate samples), and the last two columns list the mean and standard deviation of the replicate marginal-likelihood estimates. Candidate models are listed in rows and comprise 12 combinations of: (1) instantaneous-rate matrices (symmetric, $\mathbf{Q}_s$ or asymmetric, $\mathbf{Q}_a$); (2) priors on the average dispersal rate [default, $P_d(\mu)$ or alternative, $P_a(\mu)$]; (3) priors on the number of dispersal routes [default, $P_d(\Delta)$ or alternative, $P_a(\Delta)$], and; (4) the three models describing how dispersal dynamics and average dispersal rates vary through time (constant, $I_1$, two-intervals, $I_2$, or four-intervals, $I_4$). Columns 1 and 2 list the index, $M_i$, and parameterization of each candidate biogeographic model, respectively. The preferred biogeographic model is indicated in bold text.

| Index | Candidate model | replicate 1 | replicate 2 | replicate 3 | replicate 4 | combined | mean | sd |
|---|---|---|---|---|---|---|---|---|
| $M_1$ | $I_1 P_d(\mu) \mathbf{Q}_a P_d(\Delta)$ | −2527.09 | −2526.18 | −2525.69 | −2524.43 | −2525.09 | −2525.85 | 1.11 |
| $M_2$ | $I_1 P_d(\mu) \mathbf{Q}_a P_a(\Delta)$ | −2503.84 | −2504.09 | −2505.79 | −2504.27 | −2504.21 | −2504.50 | 0.88 |
| $M_3$ | $I_1 P_d(\mu) \mathbf{Q}_s P_d(\Delta)$ | −2559.29 | −2560.11 | −2561.17 | −2560.25 | −2560.40 | −2560.21 | 0.77 |
| $M_4$ | $I_1 P_d(\mu) \mathbf{Q}_s P_a(\Delta)$ | −2504.00 | −2504.09 | −2502.86 | −2503.08 | −2503.27 | −2503.51 | 0.63 |
| $M_5$ | $I_1 P_a(\mu) \mathbf{Q}_a P_d(\Delta)$ | −2166.40 | −2164.48 | −2162.31 | −2165.59 | −2164.12 | −2164.69 | 1.78 |
| $M_6$ | $I_1 P_a(\mu) \mathbf{Q}_a P_a(\Delta)$ | −2161.51 | −2160.05 | −2160.49 | −2160.21 | −2160.00 | −2160.57 | 0.65 |
| $M_7$ | $I_1 P_a(\mu) \mathbf{Q}_s P_d(\Delta)$ | −2255.39 | −2255.53 | −2260.00 | −2255.15 | −2256.40 | −2256.52 | 2.33 |
| $M_8$ | $I_1 P_a(\mu) \mathbf{Q}_s P_a(\Delta)$ | −2200.05 | −2200.87 | −2200.58 | −2200.18 | −2200.16 | −2200.42 | 0.37 |
| $M_9$ | $I_2 P_d(\mu) \mathbf{Q}_a P_d(\Delta)$ | −2591.04 | −2591.15 | −2590.85 | −2591.77 | −2590.20 | −2591.20 | 0.40 |
| $M_{10}$ | $I_4 P_d(\mu) \mathbf{Q}_a P_d(\Delta)$ | −2644.56 | −2647.48 | −2646.01 | −2642.56 | −2642.90 | −2645.15 | 2.10 |
| $M_{11}$ | $I_2 P_a(\mu) \mathbf{Q}_a P_a(\Delta)$ | −2155.95 | −2157.00 | −2155.37 | −2158.00 | −2155.98 | −2156.58 | 1.17 |
| $\boldsymbol{M_{12}}$ | $\boldsymbol{I_4 P_a(\mu) \mathbf{Q}_a P_a(\Delta)}$ | **−2138.31** | **−2138.12** | **−2136.96** | **−2136.59** | **−2136.51** | **−2137.49** | **0.85** |

**The absolute fit of competing biogeographic models to the reduced SARS-CoV-2 dataset**

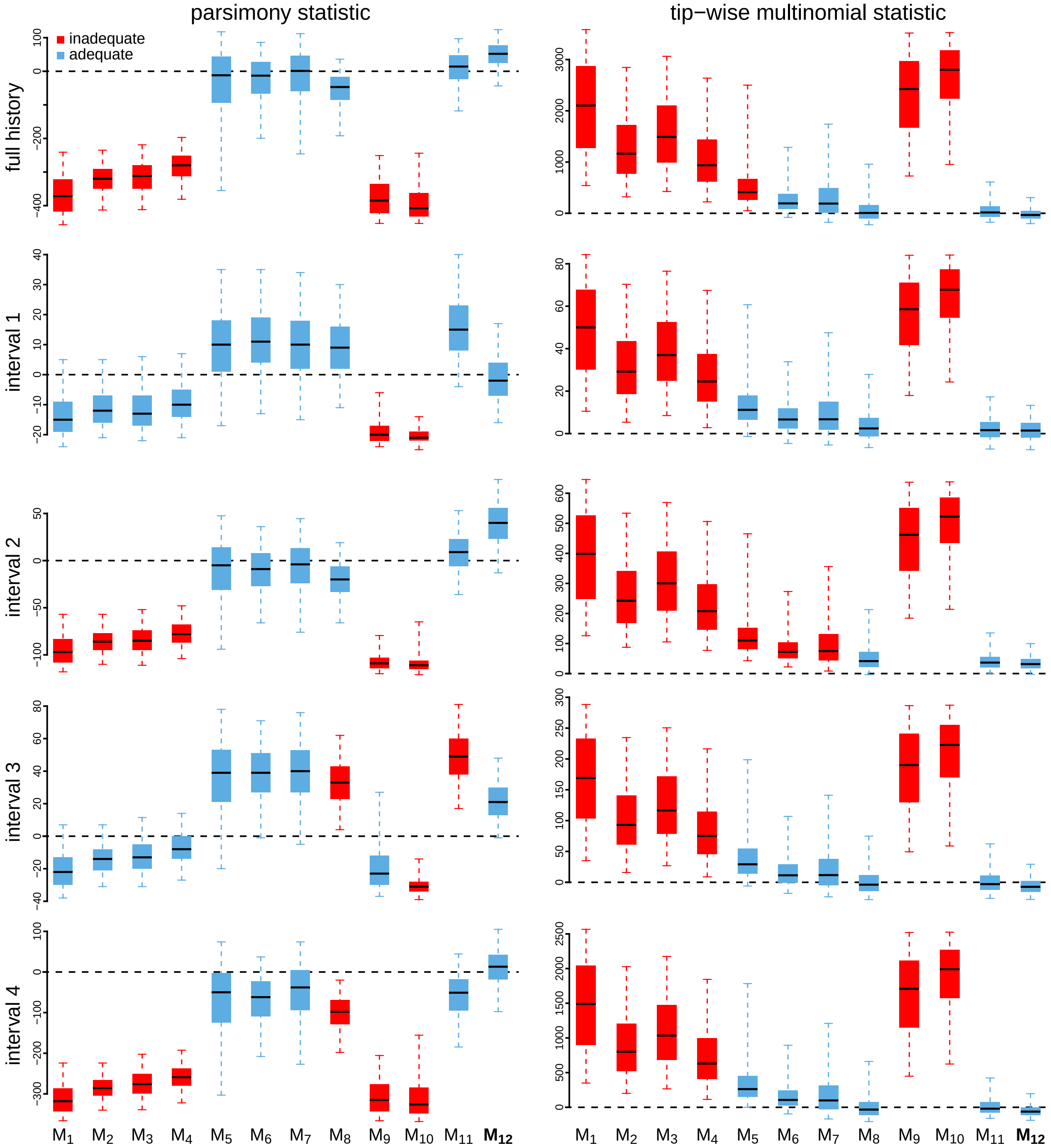


Figure S5: **Posterior-predictive distributions under the candidate biogeographic models conditioning on the MCC phylogeny for the reduced SARS-CoV-2 dataset.** The left column depicts posterior-predictive distributions for the parsimony statistic; the right column depicts distributions for the tip-wise multinomial statistic. The first row reflects statistics computed for the entire period spanning the early phase of the pandemic, while the remaining four rows depict statistics computed for each of the four time intervals specified in the preferred piecewise-constant biogeographic model. Each panel includes a set of 12 box plots (one for each candidate biogeographic model presented in Table S4). The center of each box is the median predictive value of the test statistic; the box and whiskers indicate the corresponding 50% and 95% posterior-predictive intervals, respectively. The horizontal dashed line indicates the value of the test statistic under identical fit of the simulated and observed datasets. A model is judged to be inadequate (*i.e.*, incapable of generating geographic datasets that are similar to the observed data) if its 95% posterior-predictive interval does not overlap with the dashed line. The preferred biogeographic model, $M_{12}$, is indicated in bold.

#### S2.3 Joint Analyses of the Entire SARS-CoV-2 Dataset

##### Overview

In this section, we describe the analyses we performed to infer the joint posterior probability distribution of the phylodynamic model—comprising all parameters of the component relaxed-clock and biogeographic models—from the entire SARS-CoV-2 dataset (which includes the viral genome sequences, and the geographic areas and dates of viral sampling). For these analyses, we specified a relaxed-clock model that was similar to that used to estimate the dated phylogeny for the reduced SARS-CoV-2 dataset in Section S2.1, and specified the biogeographic model that was selected based on analyses of the reduced dataset in Section S2.2. Below, we provide details on: (1) the specified phylodynamic model; (2) the MCMC simulations we performed to estimate the joint posterior under this model, and; (3) the posterior-predictive simulations we performed to assess the absolute fit of the biogeographic model to the entire SARS-CoV-2 dataset.

###### Model specification

Our joint analyses of the entire SARS-CoV-2 dataset are based on a phylodynamic model that includes (1) a relaxed-clock model, and (2) a biogeographic model. The relaxed-clock model that we specified for our joint analyses of the entire SARS-CoV-2 sequence dataset is identical to that specified previously in our analyses of the reduced SARS-CoV-2 sequence dataset (see Section S2.1) with minor changes with prior specification to accommodate differences in viral sampling (see Table S5). The biogeographic model that we specified for our joint analyses of the entire SARS-CoV-2 geographic dataset is identical to the biogeographic model that we selected previously based on analyses of the reduced SARS-CoV-2 dataset (see Section S2.2); specifically, the 4-interval piecewise-constant alternative-prior asymmetric geographic model.

###### Data analysis

*Estimating the joint posterior of phylodynamic model parameters using MCMC simulation*

We performed 20–30 independent MCMC simulations to approximate the joint posterior distribution of the phylodynamic-model parameters—including the phylogeny, divergence times and biogeographic history—from the entire SARS-CoV-2 dataset using our modified version of `BEAST` (see Section S3 for details) with the `BEAGLE` library (compiled from the 'hmc-clock' branch, commit 'dd36bf5': `https://github.com/beagle-dev/beagle-lib/tree/dd36bf5b8d88348c77a93eeeef0917d90df71a4f`; Ayres et al. 2019) enabled to accelerate computation. We ran each replicate MCMC simulation for 10–20 mil-

Table S5: **Priors used to jointly infer SARS-CoV-2 phylogeny and biogeographic history for the entire dataset.**

| Parameter | Description | Prior |
|---|---|---|
| $\kappa_1$ | Ratio of the A $\rightarrow$ G rate to the transversion rate | Lognormal$(\mu = 1.0, \sigma = 0.8)^*$ |
| $\kappa_2$ | Ratio of the C $\rightarrow$ T rate to the transversion rate | Lognormal$(\mu = 1.0, \sigma = 0.8)$ |
| $\boldsymbol{\pi}$ | Nucleotide stationary frequencies | Dir$(1,1,1,1)$ |
| $\boldsymbol{m}$ | Partition-specific rate multipliers | Dir$(1,1,1,1,1,1)$ |
| $\alpha$ | Shape and scale parameter of the $\Gamma_4$ distribution | Lognormal$(\mu = -2.1, \sigma = 0.5874)$ |
| $\mathbb{E}[\boldsymbol{r}]$ | Mean of the UCLN | Lognormal$(\mu = -12.5, \sigma = 0.5)$ |
| $SD(\boldsymbol{r})$ | Standard deviation of the UCLN | Exp$(\lambda = 1/(4.0\text{e}{-}6))$ |
| $N_T$ | Effective number of infected individuals at sampling time, $T$ | Lognormal$(\mu = 7.0, \sigma = 1.0)$ |
| $r$ | Exponential growth rate of the coalescent model | Laplace$(0.07, 0.01)$ |
| $\Delta_l$ | Number of dispersal routes in interval $l$ | Pois$(253)$ |
| $\mu_l$ | Average dispersal rate in interval $l$ | Exp$(1/\lambda)$; $\lambda \sim \Gamma(0.5, 0.5)$ |
| $r_{ij,l}$ | Relative dispersal rate from $i$ to $j$ in interval $l$ | $\Gamma(1,1)$ |

*$\mu$ and $\sigma$ in this table are the mean and standard deviation of the normal distribution.

lion generations, sampling continuous parameters every 1000 generations and trees every 10,000 generations. Details of these analyses are available in the XML scripts included in our GitHub and Dryad repositories.

After discarding the first 10–75% of samples from each replicate MCMC simulation (as burn-in), we combined the remaining posterior samples of trees from all replicates and then down-sampled every 50,000 generations using `LogCombiner` version 1.10.5. Following initial inspection of the log files using Tracer (Rambaut et al. 2018) version 1.7.1, we further evaluated MCMC performance using the `coda` package (Plummer et al. 2006) in `R` (R Core Team 2020). We assessed convergence of replicate MCMC simulations by calculating the ESS for each continuous parameter for the combined posterior samples; ensuring that values for the substitution-model parameters were all $\gg 4000$, those for the geographic model parameters were all $\gg 200$, and that the ESS values for all parameters of the branch-rate and branching-process models were all $\gg 100$.

*(Re)assessing adequacy of the biogeographic model using posterior-predictive simulation*

We previously established that the preferred biogeographic model provides an adequate description of the process of geographic dispersal during the early phase of the COVID-19 pandemic (see Section S2.2). However, those analyses were based on the reduced (rather than entire) SARS-CoV-2 dataset, and also conditioned on a single dated phylogeny—the MCC tree inferred in Section S2.1—rather than integrating over the posterior probability distribution of dated phylogenies. Accordingly, we performed analyses to confirm that the preferred biogeographic model provides an adequate fit to the entire SARS-CoV-2 dataset under an inference scenario where geographic history is jointly integrated over the posterior distribution of dated phylogenies.

We performed a series of posterior-predictive simulations to assess the adequacy of the preferred (the 4-interval piecewise-constant alternative-prior asymmetric) biogeographic model. As a point of reference, we also assessed the absolute fit of the best time-constant biogeographic model (*c.f.*, Table S4). For both biogeographic models, we simulated $m = 2500$ predictive datasets by repeatedly sampling at random from the corresponding joint posterior distribution of phylodynamic model parameters inferred from the entire SARS-CoV-2 dataset. We then generated posterior-predictive distributions from each set of $m$ predictive datasets under 10 separate test statistics, *i.e.*, for all combinations of the two (parsimony and tip-wise multinomial) summary statistics, and each of five time intervals (*i.e.*, the entire history and each of the four intervals of the piecewise-constant biogeographic model). For each posterior-predictive distribution, we computed the posterior-predictive p-value (see Section S3.3 for details).

**Results**

Posterior-predictive simulations confirm that the preferred biogeographic model (4-interval piecewise-constant alternative-prior asymmetric geographic model) provides an adequate fit to the entire SARS-CoV-2 dataset, whereas the best time-constant biogeographic model is inferred to be inadequate (Figures S6 and S9). The results of these joint analyses of the entire SARS-CoV-2 dataset are presented in the main text (Fig. 1B, C) and Fig. S7.

#### The absolute fit of the preferred biogeographic model to the entire SARS-CoV-2 dataset

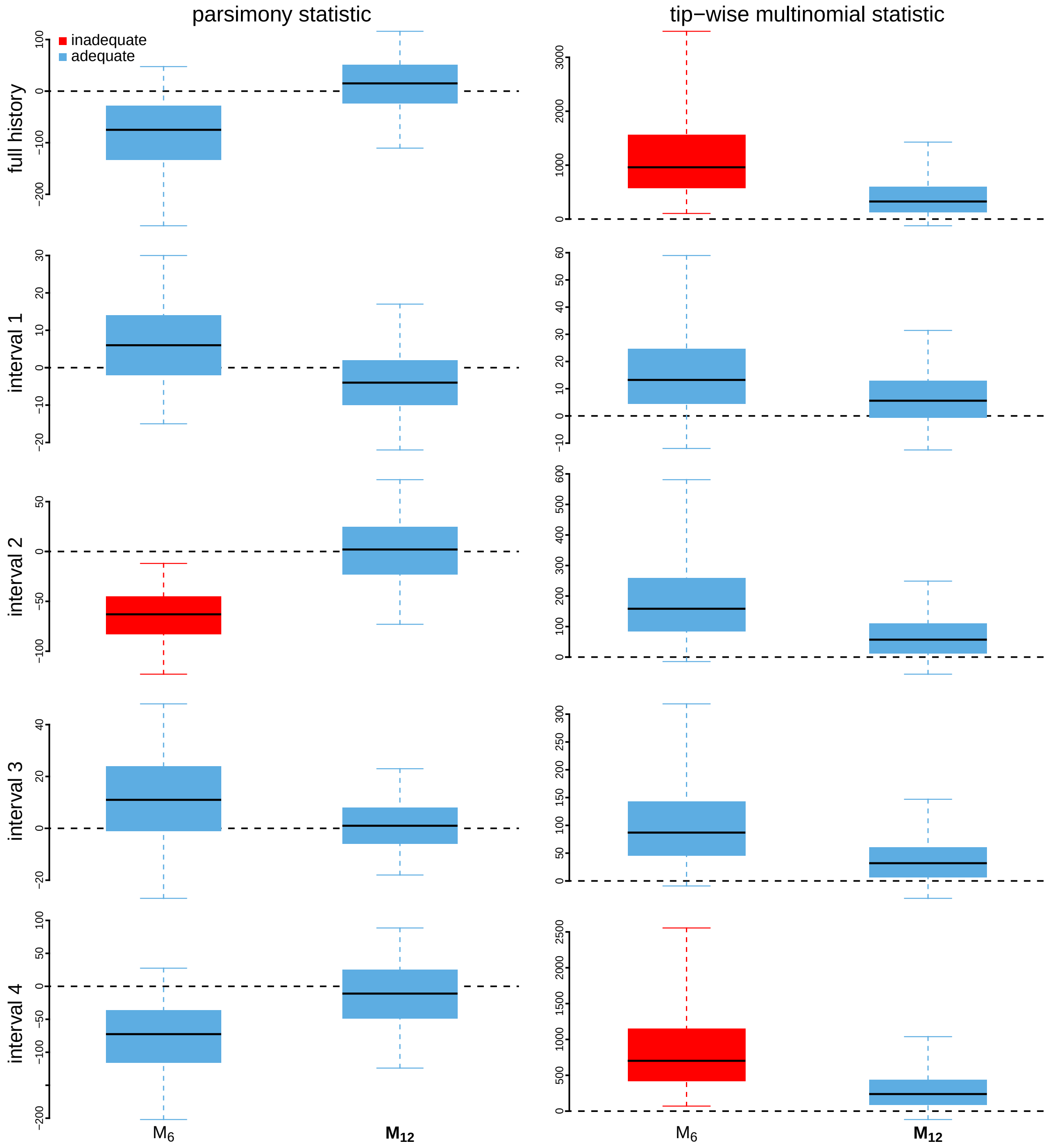


Figure S6: **Posterior-predictive distributions of biogeographic models under joint inference of the entire SARS-CoV-2 dataset.** The left column depicts posterior-predictive distributions for the parsimony statistic; the right column depicts distributions for the tip-wise multinomial statistic. The first row reflects statistics computed for the entire period spanning the early phase of the pandemic, while the remaining four rows depict statistics computed for each of the four time intervals specified in the preferred piecewise-constant biogeographic model. Each panel includes boxplots for two biogeographic models: the best time-constant model, $M_6$, and the preferred piecewise-constant model, $M_{12}$ (*c.f.*, Table S4). The center of each box is the median predictive value of the test statistic; the box and whiskers indicate the corresponding 50% and 95% posterior-predictive intervals, respectively. The horizontal dashed line indicates the value of the test statistic under identical fit of the simulated and observed datasets. A model is judged to be inadequate (*i.e.*, incapable of generating geographic datasets that are similar to the observed data) if its 95% posterior-predictive interval does not overlap with the dashed line.

#### Inferred pairwise dispersal parameters under the piecewise-constant model

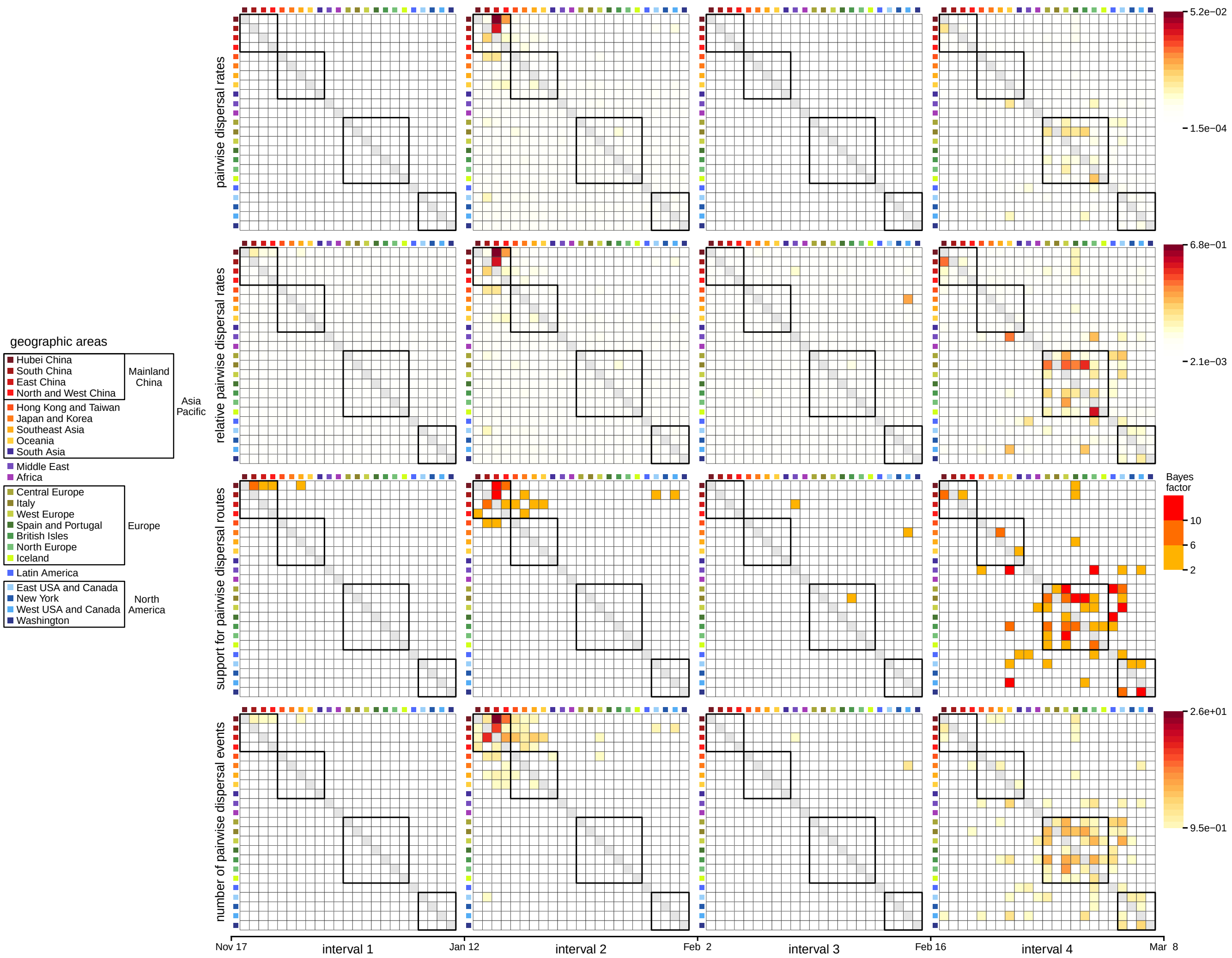


Figure S7: **Summary of dispersal parameters inferred from joint analyses of the entire SARS-CoV-2 dataset.** The four time intervals exhibit distinct dispersal dynamics. **(A)** Absolute viral dispersal rate between each pair of discrete geographic areas. **(B)** Relative viral dispersal rate (*i.e.*, the absolute rate in panel A divided by the inferred global dispersal rate for the corresponding interval) between each pair of discrete geographic areas. **(C)** The evidential support (Bayes factors, inset legend, panel C, right) that a given dispersal route between a pair of discrete geographic areas played a role in the spread of the virus. **(D)** Number of viral dispersal events between each pair of discrete geographic areas. Boxes in each panel indicate groups of areas (inset legend, left). The first interval is dominated by dispersal from Hubei to other areas in China, the second interval by more widespread dispersal within Asia and by dispersal from China to North America, culminating in cosmopolitan dispersal in the fourth interval. Note that interval three—immediately following the onset of international air-travel bans with China—exhibits a reduction in the number of viral dispersal routes, including disruption of the dispersal routes from China to North America.

#### S2.4 Estimating Daily Global Viral Dispersal Rates

##### Overview

In this section, we describe our analyses to explore the correlation between daily global air-travel volume and global SARS-CoV-2 dispersal rates during the early phase of the COVID-19 pandemic. Because we were able to obtain data on the *daily* volume of global air travel during this period, we performed an analysis of the entire SARS-CoV-2 dataset under a more granular phylodynamic model that allows the average dispersal rate to vary from day to day. We used these analyses to infer the number of daily viral dispersal events between pairs of geographic areas. We then performed standard statistical tests to assess the degree of correlation between the inferred daily global viral dispersal rates and daily global air-travel volume.

##### Model specification

Our estimates of daily variation in global viral dispersal rates are based on a phylodynamic model that is identical to the model we previously used to infer the joint posterior of SARS-CoV-2 phylogeny and biogeographic history (see Section S2.3), except that we further discretized the number of time intervals in which the global dispersal rate, $\mu$, was free to vary. Specifically, rather than allowing global dispersal rates to vary between four time intervals, we specified an independent $\mu$ for each day between December 30, 2019 and March 8, 2020 (with an additional independent $\mu$ for the time spanning the origin of SARS-CoV-2 to December 29, 2019). The prior on each $\mu$ is specified according to the posterior estimates inferred in Section S2.3. We computed the posterior mean of the global viral dispersal rate across the entire history inferred from the joint analyses and used it as the prior mean, and we specified standard deviation of the prior distribution so that the 95% prior interval spans three orders of magnitude around the mean. Details of the priors are described in Table S6. Our inferences of geographic history under this model were averaged over the marginal posterior probability distribution of dated phylogenies inferred in Section S2.3. Details of these analyses are available in the XML scripts included in our GitHub and Dryad repositories.

##### Data analysis

###### Parameter estimation

We performed 15 independent MCMC simulations to approximate the joint posterior distribution of the biogeographic-model parameters from the entire SARS-CoV-2 dataset using our modified version of `BEAST` (see Section S3 for details) and `BEAGLE` version 3.2.0 (Ayres et al. 2019). We ran each replicate MCMC simulation for 5 million generations, sampling continuous parameters every 2000 generations and phylogenies every 10000 generations. Every 10000 generations we simulated full dispersal histories over the tree sampled in that generation. Details of these analyses are available in the XML scripts included in our GitHub and Dryad repositories.

For each replicate MCMC simulation, we discarded the first million generations (as burn-in), and then combined the remaining posterior samples from all replicates using `LogCombiner` version 1.10.5. Following initial inspection of the log files using Tracer (Rambaut et al. 2018) version 1.7.1, we further

Table S6: **Priors used to infer the daily global viral dispersal rates for the entire dataset.**

| Parameter | Description | Prior |
|---|---|---|
| $\Delta_l$ | Number of dispersal routes in interval $l$ | Pois(253) |
| $\mu_p$ | Global dispersal rate in day $p$ | $\text{Lognormal}(\mu = -4.76, \sigma = 1.7622)^*$ |
| $r_{ij,l}$ | Relative dispersal rate from $i$ to $j$ in interval $l$ | $\Gamma(1, 1)$ |

$^*\mu$ and $\sigma$ in this table are the mean and standard deviation of the normal distribution.

evaluated MCMC performance using the `coda` package (Plummer et al. 2006) in `R` (R Core Team 2020). We assessed convergence of replicate MCMC simulations by calculating the ESS for each continuous parameter for the combined posterior samples, ensuring that values for all parameters were $\gg 700$.

##### Correlation test

We performed a standard test for correlation between the volume of daily global air travel and the rate of daily global SARS-CoV-2 dispersal by computing Pearson's $r$ and the corresponding $p$ value. We focussed on the period spanning from January 31—when the virus first achieved a cosmopolitan distribution, WHO (2020)—to March 8, 2020.

##### Parameter summary

We summarized the number of viral dispersal events between a given pair of geographic areas by counting the number of dispersal events from the source region (*e.g.*, China) to the destination region (*e.g.*, North America) that occurred on that day for a given simulated history, and then looped over all of the histories to obtain the posterior distribution of the number of pairwise dispersal events. Mean and 95% credible intervals for the daily number of viral dispersal events were then computed from the corresponding posterior distribution.

#### Results

The results of these analyses are presented in the main text (see Figs. 1B and 2A), as well as Fig. S8 below.

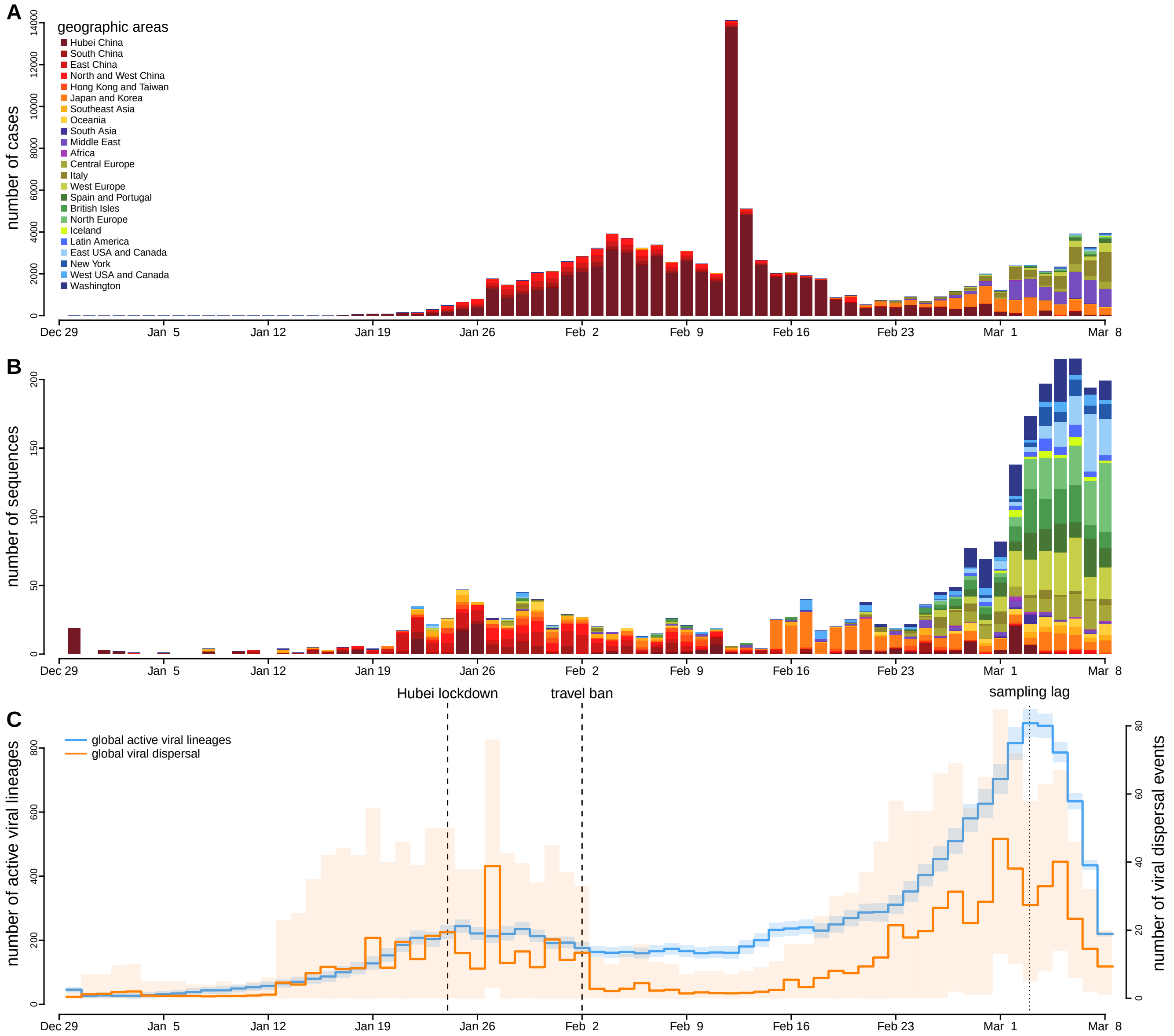


Figure S8: **Global prevalence of COVID-19 and the inferred number of SARS-CoV-2 dispersal events. (A)** Daily number of COVID-19 confirmed cases during the early phase of the pandemic. **(B)** Daily number of sampled SARS-CoV-2 sequences in the entire dataset. Stacked-bar plot colors in panels A and B correspond to the areas where cases were reported/viruses sampled (legend, panel A). **(C)** Inferred number of active SARS-CoV-2 lineages (posterior mean [blue solid lines], 95% credible interval [blue shaded areas]) and number of viral dispersal events (posterior mean [orange solid lines], 95% credible interval [orange shaded areas]). A sharp decrease in the inferred number of global viral dispersal events on February 2—which coincides with the initiation of the international travel bans—is due to the decrease of the global viral dispersal rate (Fig. 1B, main text). The number of global viral dispersal events increases gradually after February 16 as a result of the increasing global viral dispersal rate (Fig. 1B, main text) and also the increasing number of actively circulating viral lineages. Note that sampling lag causes the number of viral dispersal events close to the end of the sampling period to be underestimated. Unsurprisingly, the inferred number of active viral lineages more closely matches the number of sampled SARS-CoV-2 sequences in our dataset than the number of confirmed COVID-19 cases.

#### S2.5 Assessing the Impact of Incomplete Viral Sampling

##### Overview

The results of our phylodynamic analyses are based on a small and non-random sample of the total population of SARS-CoV-2 viruses that existed during the early phase of the COVID-19 pandemic. In this section, we describe an additional series of analyses that we performed to assess the sensitivity of our results to incomplete and non-random sampling sampling of SARS-CoV-2 sequences. Specifically, we replicated the entire series of analyses that we performed on the entire SARS-CoV-2 dataset—including joint inference of phylogeny, divergence times and biogeographic history (see Section S2.3), as well as analyses to infer daily global viral dispersal rates (see Section S2.4)—for the reduced SARS-CoV-2 dataset, which has fewer (1271 vs 2598) sequences and different temporal and spatial sampling intensities.

##### Model specification

###### Joint phylodynamic inference

Our joint analyses of the reduced SARS-CoV-2 dataset are based on a phylodynamic model that includes (1) a relaxed-clock model, and (2) a biogeographic model. The relaxed-clock model that we specified for our joint analyses of the reduced SARS-CoV-2 dataset is identical to that specified previously in our analyses to estimate a dated phylogeny for this sequence dataset (see Section S2.1). The biogeographic model that we specified for our joint phylodynamic analyses of the reduced SARS-CoV-2 dataset is identical to the biogeographic model that we selected previously based on analyses of this geographic dataset that conditioned on the MCC summary tree (see Section S2.2); specifically, the 4-interval piecewise-constant alternative-prior asymmetric geographic model. Details of the priors used in these analyses are described in Table S7.

###### Estimating daily global viral dispersal rates

Our estimates of daily variation in global viral dispersal rates are based on a phylodynamic model that is identical to the model we used to infer the joint posterior of SARS-CoV-2 phylogeny and biogeographic history, except that we further discretized the number of time intervals in which the global dispersal rate, $\mu$, was free to vary. Specifically, rather than allowing global dispersal rates to vary between four time intervals, we specified an independent $\mu$ for each day between December 30, 2019 and March 8, 2020 (with an additional independent $\mu$ for the time spanning the origin of SARS-CoV-2 to December 29, 2019). The prior on each $\mu$ is specified according to the posterior estimates inferred in Section S2.3.

Table S7: **Priors used to jointly infer SARS-CoV-2 phylogeny and dispersal history for the reduced dataset.**

| Parameter | Description | Prior |
|---|---|---|
| $\kappa_1$ | Ratio of the A $\rightarrow$ G rate to the transversion rate | Lognormal$(\mu = 1.0, \sigma = 0.8)^*$ |
| $\kappa_2$ | Ratio of the C $\rightarrow$ T rate to the transversion rate | Lognormal$(\mu = 1.0, \sigma = 0.8)$ |
| $\boldsymbol{\pi}$ | Nucleotide stationary frequencies | Dir$(1,1,1,1)$ |
| $\boldsymbol{m}$ | Partition-specific rate multipliers | Dir$(1,1,1,1,1,1)$ |
| $\alpha$ | Shape and scale parameter of the $\Gamma_4$ distribution | Lognormal$(\mu = -2.1, \sigma = 0.5874)$ |
| $\mathbb{E}[\boldsymbol{r}]$ | Mean of the UCLN | Lognormal$(\mu = -12.7, \sigma = 0.5874)$ |
| $SD(\boldsymbol{r})$ | Standard deviation of the UCLN | Exp$(\lambda = 1/(2.0\text{e}{-6}))$ |
| $N_T$ | Effective number of infected individuals at sampling time, $T$ | Lognormal$(\mu = 7.5, \sigma = 1.0)$ |
| $r$ | Exponential growth rate of the coalescent model | Laplace$(0.07, 0.01)$ |
| $\Delta_l$ | Number of dispersal routes in interval $l$ | Pois$(253)$ |
| $\mu_l$ | Average dispersal rate in interval $l$ | Exp$(1/\lambda)$; $\lambda \sim \Gamma(0.5, 0.5)$ |
| $r_{ij,l}$ | Relative dispersal rate from $i$ to $j$ in interval $l$ | $\Gamma(1,1)$ |

$^*\mu$ and $\sigma$ in this table are the mean and standard deviation of the normal distribution.

Table S8: **Priors used to infer the daily global viral dispersal rates for the reduced dataset.**

| Parameter | Description | Prior |
|---|---|---|
| $\Delta_l$ | Number of dispersal routes in interval $l$ | $\text{Pois}(253)$ |
| $\mu_p$ | Global dispersal rate in day $p$ | $\text{Lognormal}(\mu = -4.28, \sigma = 1.7622)^*$ |
| $r_{ij,l}$ | Relative dispersal rate from $i$ to $j$ in interval $l$ | $\Gamma(1, 1)$ |

*$\mu$ and $\sigma$ in this table are the mean and standard deviation of the normal distribution.

We computed the posterior mean of the global viral dispersal rate across the entire history inferred from the joint analyses and used it as the prior mean, and we specified standard deviation of the prior distribution so that the 95% prior interval spans three orders of magnitude around the mean. Details of the priors used in these analyses are described in Table S8.

#### Data Analysis

##### Estimating the joint posterior of phylodynamic model parameters using MCMC simulation

Our analyses to estimate the joint posterior probability distribution of the phylodynamic-model parameters—including the phylogeny, divergence times and biogeographic history—from the reduced SARS-CoV-2 dataset were identical to those used in our joint analyses of the entire SARS-CoV-2 dataset (see Section S2.3). Details of these analyses are available in the XML scripts included in our GitHub and Dryad repositories.

##### (Re)assessing adequacy of the biogeographic model using posterior-predictive simulation

We previously established that the preferred biogeographic model provides an adequate description of the process of geographic dispersal that gave rise to the reduced SARS-CoV-2 dataset during the early phase of the COVID-19 pandemic (see Section S2.2). However, those analyses were conditioned on a single dated phylogeny—the MCC tree inferred in Section S2.1—rather than integrating over the posterior probability distribution of dated phylogenies. Accordingly, we performed analyses to confirm that the preferred biogeographic model provides an adequate fit to the reduced SARS-CoV-2 dataset under an inference scenario where geographic history is jointly integrated over the posterior distribution of dated phylogenies.

We performed a series of posterior-predictive simulations to assess the adequacy of the preferred, 4-interval piecewise-constant alternative-prior asymmetric biogeographic model, $M_{12}$ (*c.f.*, Table S4). For context, we also assessed the absolute fit of the best time-constant biogeographic model, $M_6$, and the best 2-interval piecewise-constant biogeographic model, $M_{11}$. All other aspects of these posterior-predictive simulations of the reduced SARS-CoV-2 dataset were identical to those used in our joint analyses of the entire SARS-CoV-2 dataset (see Section S2.3).

##### Estimating daily number of dispersal events and global viral dispersal rates

We performed analyses of the reduced SARS-CoV-2 dataset to infer daily global viral dispersal rates and estimate the number of daily number of pairwise dispersal events of the reduced SARS-CoV-2 dataset. These analyses are identical to those performed on the entire SARS-CoV-2 dataset (see Section S2.4).

#### Results

Posterior-predictive simulations confirm that the preferred (4-interval piecewise-constant alternative-prior asymmetric) biogeographic model provides an adequate fit to the reduced SARS-CoV-2 dataset under a joint-inference scenario (Figure S9). Importantly, results based on our analyses of the reduced SARS-CoV-2 dataset (Figs. S10, S11, S12) are qualitatively identical to those based on analyses of the entire dataset (Figs. 1B, 1C, 2A), suggesting that our findings are robust to sampling artifacts.

**The absolute fit of competing biogeographic models to the reduced SARS-CoV-2 dataset**

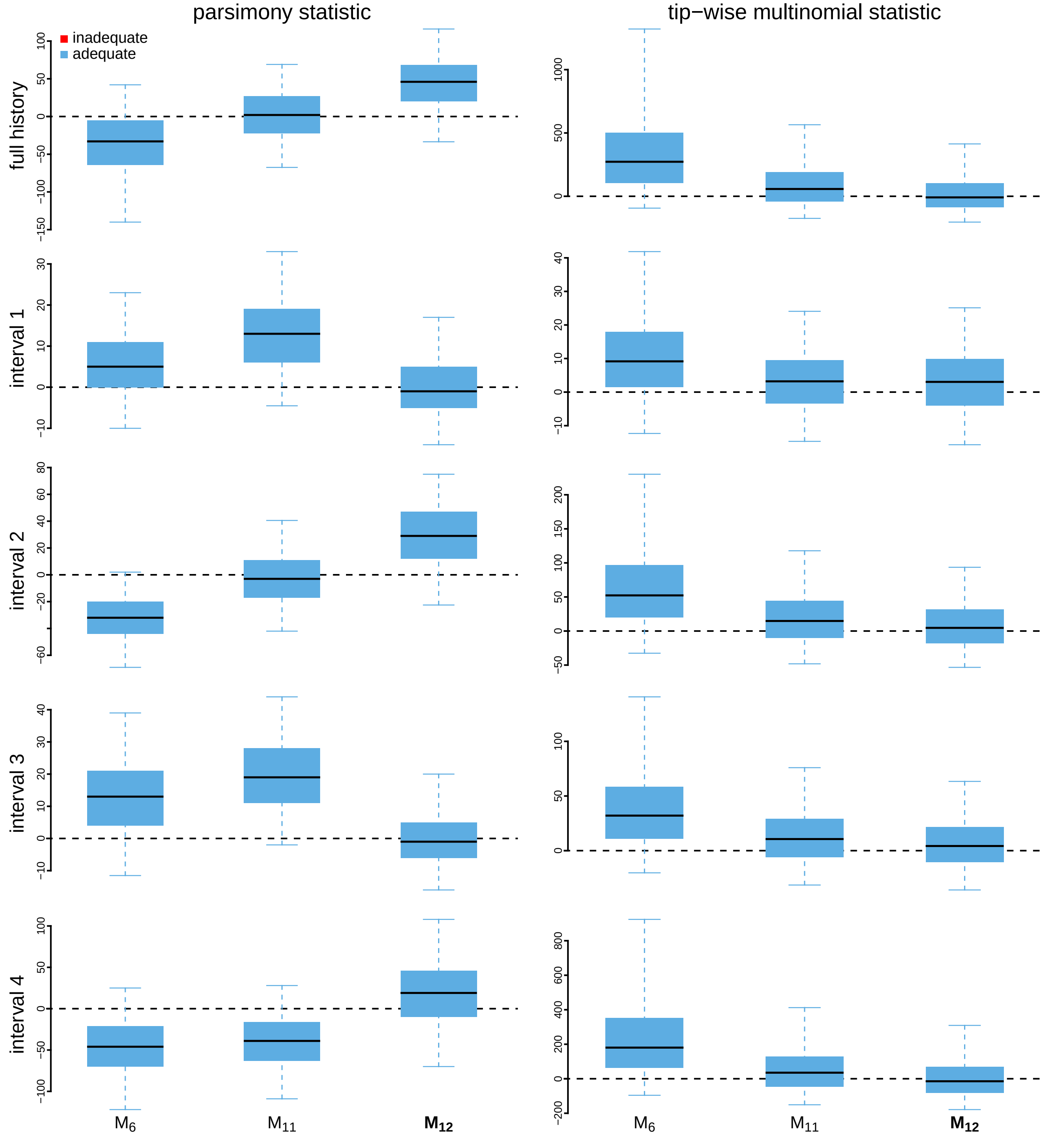


Figure S9: **Posterior-predictive distributions under the candidate biogeographic models based on joint inferene of the reduced SARS-CoV-2 dataset.** The left column depicts posterior-predictive distributions for the parsimony statistic; the right column depicts distributions for the tip-wise multinomial statistic. The first row reflects statistics computed for the entire period spanning the early phase of the pandemic, while the remaining four rows depict statistics computed for each of the four time intervals specified in the preferred piecewise-constant biogeographic model. Each panel includes boxplots for three biogeographic models: the best time-constant model, $M_6$, the best 2-interval piecewise-constant biogeographic model, $M_{11}$, and the preferred piecewise-constant model, $M_{12}$ (*c.f.*, Table S4). The center of each box is the median predictive value of the test statistic; the box and whiskers indicate the corresponding 50% and 95% posterior-predictive intervals, respectively. The horizontal dashed line indicates the value of the test statistic under identical fit of the simulated and observed datasets. A model is judged to be inadequate (*i.e.*, incapable of generating geographic datasets that are similar to the observed data) if its 95% posterior-predictive interval does not overlap with the dashed line.

#### Pairwise dispersal parameters inferred from the reduced SARS-CoV-2 dataset under the preferred biogeographic model

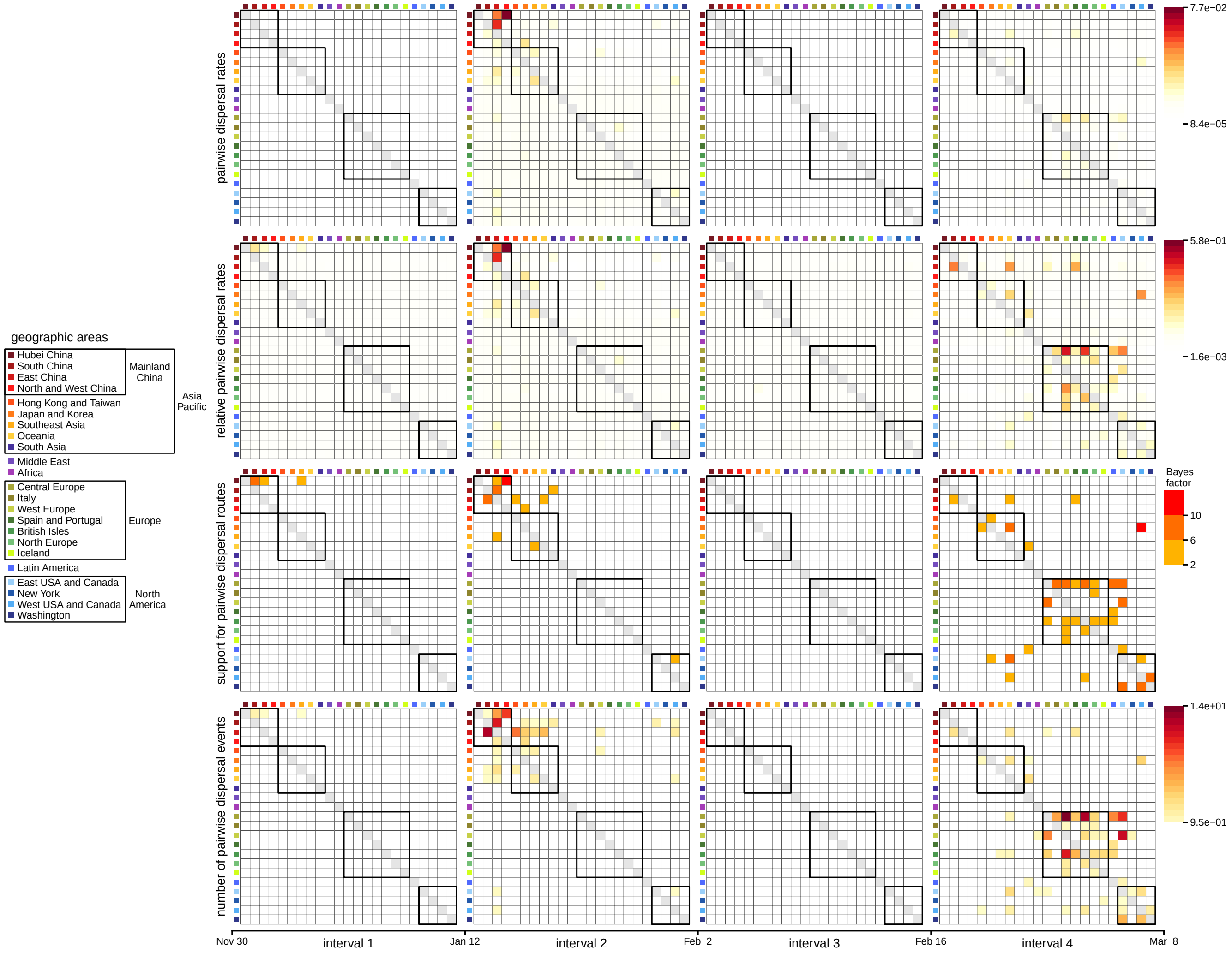


Figure S10: **Summary of pairwise dispersal parameters inferred from joint analyses of the reduced SARS-CoV-2 dataset.** The four time intervals exhibit distinct dispersal dynamics. **(A)** Absolute viral dispersal rate between each pair of discrete geographic areas. **(B)** Relative viral dispersal rate (*i.e.*, the absolute rate in panel A divided by the inferred global dispersal rate for the corresponding interval) between each pair of discrete geographic areas. **(C)** The evidential support (Bayes factors, inset legend, panel C, right) that a given dispersal route between a pair of discrete geographic areas played a role in the spread of the virus. **(D)** Number of viral dispersal events between each pair of discrete geographic areas. Boxes in each panel indicate groups of areas (inset legend, left). The first interval is dominated by dispersal from Hubei to other areas in China, the second interval by more widespread dispersal within Asia and by dispersal from China to North America, culminating in cosmopolitan dispersal in the fourth interval. Note that interval three—immediately following the onset of international air-travel bans with China—exhibits a reduction in the number of viral dispersal routes, including disruption of the dispersal routes from China to North America.

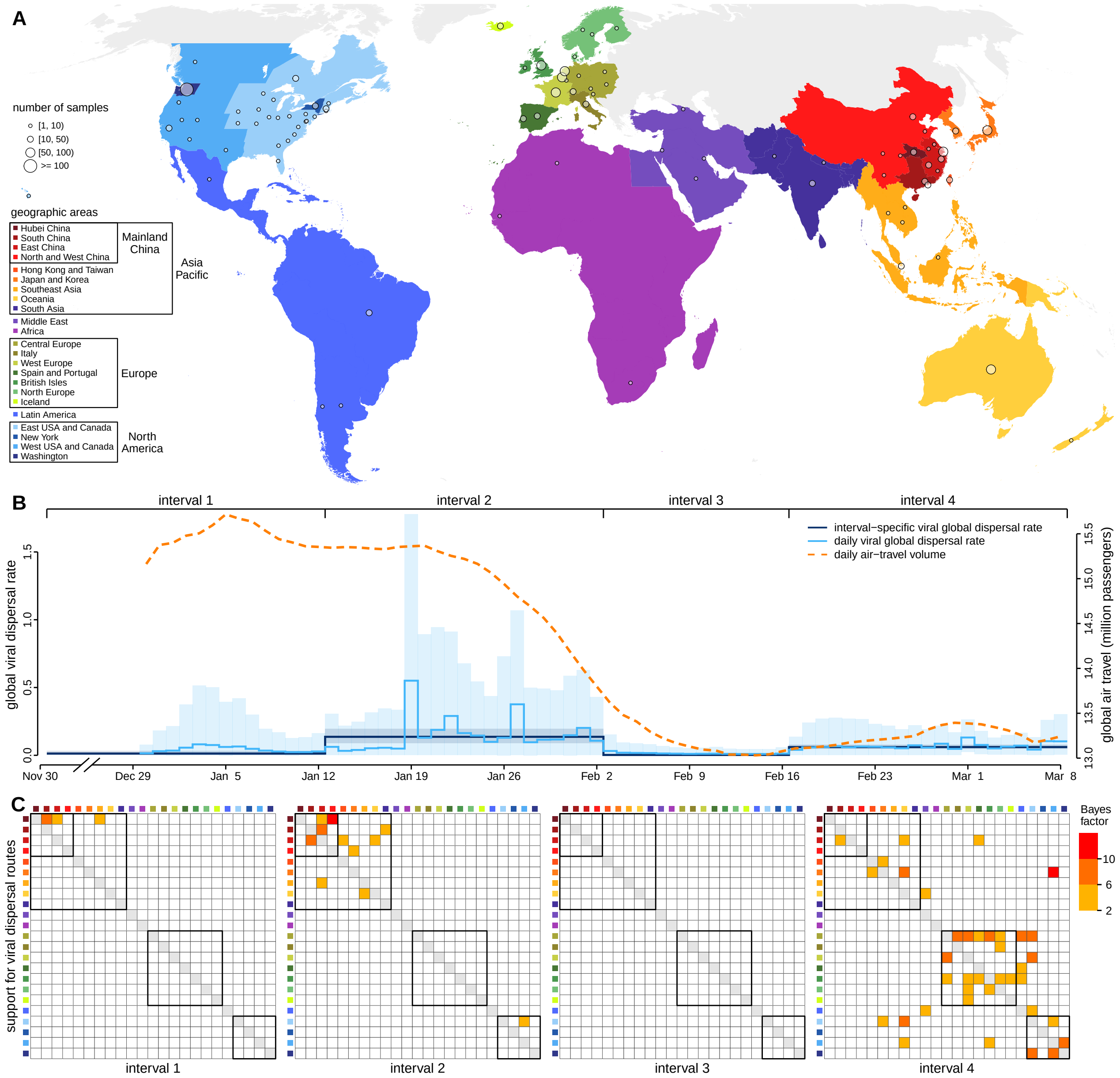


Figure S11: **Phylodynamic analyses of the reduced SARS-CoV-2 dataset reveal temporal variation in rates and dynamics of viral dispersal during the early phase of the COVID-19 pandemic. (A)** Our study includes a total of 1271 viral genomes collected between Dec. 24, 2019–Mar. 8, 2020 from 23 discrete geographic areas (colored regions); circles indicate the number and location of samples in our study. **(B)** The early phase of the COVID-19 pandemic is characterized by four time intervals spanning the MRCA of our sampled viruses (Nov. 30) to the end of our sampling period (Mar. 8). Each interval exhibits substantial differences in the global viral dispersal rate, *i.e.*, the average rate of viral dispersal across all 23 areas (posterior mean [dark blue line], 95% credible interval [dark blue shaded area]). Notably, the global viral dispersal rate decreases sharply on Feb. 2, which coincides with the onset of international air-travel restrictions with China. More granular, daily estimates of the global viral dispersal rate (light blue) and independent data on global air-travel volume (the 7-day smoothed average number of passengers, dashed orange line) are significantly correlated. **(C)** The four time intervals also exhibit distinct dispersal dynamics. Heat maps indicate the level of evidential support (Bayes factors, inset legend, right) that a given dispersal route between a pair of geographic areas played a role in the spread of the virus. Boxes in each panel indicate groups of areas (inset legend, panel A). The first interval (Nov. 30–Jan. 12) is dominated by dispersal from Hubei to other areas in China, the second interval (Jan. 12–Feb. 2) exhibits more widespread dispersal within Asia and dispersal from China to North America, culminating in cosmopolitan dispersal in the fourth interval (Feb. 16–Mar. 8). The third interval (Feb. 2–Feb. 16)—immediately following the onset of international air-travel bans with China—exhibits a reduction in the number of viral dispersal routes, including disruption of the dispersal routes from China to North America.

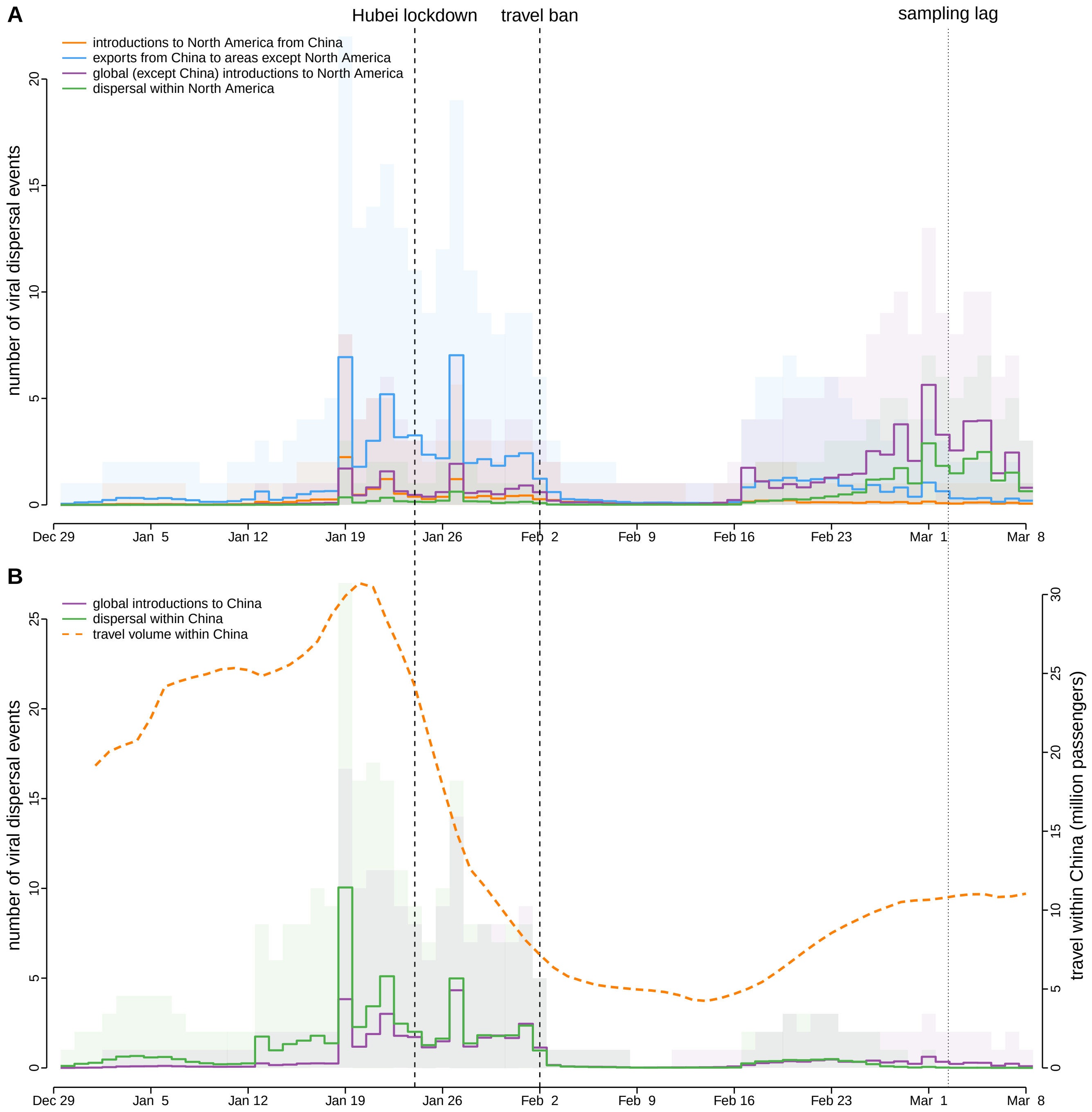


Figure S12: **The impact of COVID-19 intervention measures adopted in North America and China inferred from the reduced SARS-CoV-2 dataset.** Each panel depicts the number of viral dispersal events between areas (posterior mean [solid lines], 95% credible interval [shaded areas]). **(A)** Prior to Feb. 2, China was the major source of viral introductions into North America (orange), which explains the efficacy of the air-travel ban imposed on China. Although the travel ban had a rapid effect, it was nevertheless temporary, as—by this time—many episodes of viral dispersal from China to other regions of the world had already occurred (blue). Accordingly, the virus continued to disperse into North America from regions outside of China (purple), and also to disperse among areas within North America (green). **(B)** The partial but temporary success of the North American containment measure contrasts with the long-term impact of mitigation strategies in China, which initiated a lockdown of the Hubei province on Jan. 24, and nation-wide travel restrictions by Jan. 26. These measures drastically reduced domestic travel within China (orange dashed line), resulting in a rapid, long-term reduction in the number of viral dispersal events within China (green), despite successive waves of international viral introductions (purple). Note that sampling lag causes the number of dispersal events close to the end of the sampling period to be underestimated.

### S3 Extending Phylodynamic Methods

#### S3.1 Allowing both global dispersal rate and dispersal dynamics to vary under piecewise-constant model

Existing phylodynamic models either allow dispersal dynamics to vary over two or more pre-specified intervals (Bielejec et al. 2014)—*i.e.*, where each interval has an independent instantaneous-rate matrix, $\mathbf{Q}$, but constrains the average dispersal rate to be constant across intervals—or allow the average dispersal rate, $\mu$, to vary over two or more pre-specified intervals (Membrebe et al. 2019) (but assumes a common instantaneous-rate matrix that constrains dispersal dynamics to be constant across intervals).

Here, we extend piecewise-constant phylodynamic models to allow both the dispersal dynamics and average dispersal rate to vary independently across two or more pre-defined intervals. These models differ in how they compute the transition-probability matrix, $\mathbf{P}$, a matrix that describes the probability of transitioning from state $i$ to state $j$ over a fixed time interval, which may span changes in dispersal rates and/or dispersal dynamics.

##### Dispersal-rate variation

Under a time-constant biogeographic model, the transition-probability matrix for a branch is $\mathbf{P} = \exp(\mathbf{Q}v)$, where $v = \mu t$ represents the expected number of dispersal events on the branch with length $t$ time units and dispersal rate $\mu$. However, under the piecewise-constant model of dispersal-rate variation—where the global average dispersal rate may vary among time intervals (Membrebe et al. 2019), but the dispersal dynamics (defined by the matrix $\mathbf{Q}$) are constant across all intervals—a given branch may span two or more intervals with different average dispersal rates ("rate intervals"). In this case, the transition-probability matrix for the branch is computed as the matrix exponential:

$$\mathbf{P} = \exp(\mathbf{Q} \sum_{l=1}^{n} v_l), \tag{S1}$$

where $\mathbf{Q}$ is the instantaneous-rate matrix, $n$ is the number of rate intervals spanned by the branch, and $v_l$ is the expected number of dispersal events in interval $l$. Recall that $v_l = \mu_l t_l$, where $\mu_l$ is the average dispersal rate in interval $l$ and $t_l$ is the time spent in interval $l$.

##### Dispersal-dynamic variation

Under a piecewise-constant model of dispersal-dynamic variation—where the instantaneous-rate of dispersal between each pair of areas may vary among time intervals (Bielejec et al. 2014), but the average dispersal rate is constant across all intervals—a given branch may span two or more time intervals with different dispersal dynamics ("dynamic intervals"). In this case, the transition-probability matrix for each dynamic interval $l$, $\mathbf{P}_l$, is computed as:

$$\mathbf{P}_l = \exp(\mathbf{Q}_l v_l), \tag{S2}$$

where $\mathbf{Q}_l$ is the instantaneous-rate matrix in dynamic interval $l$, and $v_l = \mu t_l$ is the global dispersal rate multiplied by the duration of the interval. Then the transition-probability matrix for the entire branch is computed as the matrix-product of interval-wise transition-probability matrices:

$$\mathbf{P} = \prod_{l=1}^{m} \mathbf{P}_l, \tag{S3}$$

where $m$ is the number of dynamic intervals spanned by the branch.

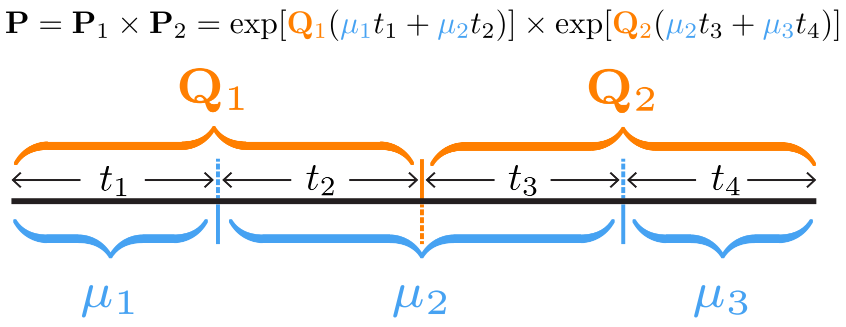


Figure S13: **Computing the transition-probability matrix for a branch spanning time intervals where both the global dispersal rate and dispersal dynamics vary.** An example illustrating the transition-probability matrix computation for a branch spanning two different dispersal dynamics ($\mathbf{Q}_1$ and $\mathbf{Q}_2$) and three different global dispersal rates ($\mu_1, \mu_2, \mu_3$).

##### Combining dispersal-rate and dispersal-dynamic variation

We combine the two approaches described above to compute transition-probability matrices under a piecewise-constant model that allows both the global dispersal rate and dispersal dynamics to vary among time intervals. Let a given branch span $m$ dynamic intervals. The expected number of dispersal events in each such dynamic interval $l$, $v_l$, is computed as:

$$v_l = \sum_{p=1}^{n} \mu_p t_p, \tag{S4}$$

where $n$ is the number of rate intervals spanned by dynamic interval $l$, $\mu_p$ is the dispersal rate in rate interval $p$, and $t_p$ is the time spent in rate interval $p$. We then substitute equation (S4) into equation (S2), and apply equation (S3) as normal to compute the transition-probability matrix for the entire branch. An example computation is illustrated in Fig. S13 for a scenario in which a branch spans two different dispersal dynamics and three different global dispersal rates.

We modified `BEAST` source code to implement the above equation for computing $\mathbf{P}$ under the piecewise-constant model that allows both $\mu$ and $\mathbf{Q}$ to vary among time intervals. An executable `BEAST` program with our extension is available in the GitHub and Dryad repositories.

#### S3.2 Stochastic mapping of geographic histories under the piecewise-constant model

Stochastic mapping, initially proposed by Nielsen (2002; see also Huelsenbeck et al. 2003; Bollback 2006; Minin and Suchard 2008), is an approach commonly used to sample viral dispersal histories over branches of a phylogeny. `BEAST` implements the endpoint-conditioned uniformization algorithm developed by Hobolth and Stone (2009) to simulate the dispersal history over each branch under the time-constant dispersal model. We extended this algorithm to sample the dispersal history under the piecewise-constant model.

Let a given branch start at time $T_0$ with state $i$ and end at time $T_m$ with state $k$. Further, let the dispersal process change (either by changing the global average dispersal rate or the dispersal dynamics) $m-1$ times on the branch at times $\{T_1, \ldots, T_{m-1}\}$, resulting in $m$ time intervals. For interval $l$, denote the dispersal rate as $\mu_l$, the instantaneous-rate matrix as $\mathbf{Q}_l$, and the duration as $t_l$. We simulate a history on this branch in a two-step procedure: first, we sample the geographic state at each time point; second, we simulate the history between each time point, conditional on the states sampled in the first step.

First, we sample geographic states at each time point. We begin by computing a transition-probability matrix for each time interval:

$$\mathbf{P}_l = \exp\left(\mathbf{Q}_l \mu_l t_l\right).$$

We then calculate the probability of state $j$ at the first time point, $T_1$, given that the branch begins in state $i$ and ends in state $k$, as:

$$\begin{aligned} \text{conditional probability } j &= \frac{\text{joint probability of } i, j, k}{\text{marginal probability of } i \text{ to } k \text{ transition}} \\ &\propto \mathbf{P}_{ij,1} \times \left[ \prod_{l=2}^{m} \mathbf{P}_l \right]_{jk}, \end{aligned}$$

where the first term is the probability of transitioning from state $i$ (the state at the beginning of the branch) to state $j$ at the first time point, and the second term is the probability of transitioning from state $j$ to state $k$ (the state at the end of the branch) over the remaining time intervals. We compute this for each state $j$, and sample the state in proportion to these probabilities. We then repeat this process for each remaining time point, recursively conditioning on the sampled state at the previous time point and the state at the end of the branch.

Second, we simulate histories within each time interval. For a given time interval, we simulate histories conditional on the start- and end-states generated in the first step using the uniformization algorithm described by Hobolth and Stone (2009).

We implemented this stochastic-mapping function both in BEAST (available in the executable included in the supplementary repositories as well) and also in our R scripts as an independent validation of the implementation (which are available in the GitHub and Dryad repositories). These two independent implementations produce effectively identical estimates of the number of viral dispersal events (Fig. S14).

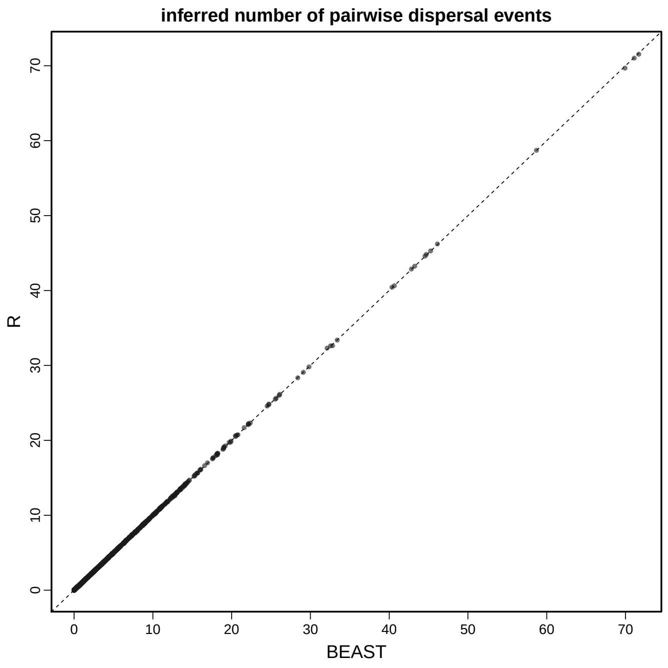


Figure S14: **Comparison of the mean estimate of the number of pairwise dispersal events in two independent implementations.** Each dot represents the mean estimate of the number of dispersal events in a given time interval between a given pair of areas. Independent implementations in BEAST and R produce effectively identical estimates.

#### S3.3 Posterior-predictive simulation for geographic models

We use posterior-predictive simulation to assess the adequacy of our biogeographic models (Gelman et al. 1996; Bollback 2002). Posterior-predictive simulation is based on the following premise: if a given model provides an adequate description of the true process that gave rise to our observed data, then datasets simulated under that model should resemble our observed dataset. In practice, we draw $m$ random samples from the joint posterior distribution of the model; each sample $i$ consists of a fully specified biogeographic model, $\theta_i = \{\Psi_i, \boldsymbol{r}_i, \boldsymbol{\delta}_i, \mu_i\}$. For each sample, we simulate a new biogeographic dataset on the sampled tree, $\Psi_i$, given the sampled parameters of the biogeographic model, $\{\boldsymbol{r}_i, \boldsymbol{\delta}_i, \mu_i\}$; we label the newly simulated dataset $G_i^{\text{sim}}$.

Next, we define a summary statistic, which we generically denote $T(G \mid \theta_i)$, where $G$ is either the simulated or observed dataset. (We note that the dependence of the statistic on $\theta_i$—while often suppressed or ignored in phylogenetic applications of posterior-predictive simulation—is consistent with posterior-predictive discrepancy analysis, as described by Gelman et al. 1996). For each simulated dataset, we compute a discrepancy statistic,

$$D_i = T(G_i^{\text{sim}} \mid \theta_i) - T(G^{\text{obs}} \mid \theta_i),$$

where $G^{\text{obs}}$ is the observed biogeographic dataset.

If the inference model provides an adequate description of the true data-generating process, the posterior-predictive distribution of $D$ should contain zero with high probability. Accordingly, for the $m$ predictive datasets for a given model and dataset combination, we calculate the posterior-predictive $p$ value as:

$$P = \frac{1}{m} \sum_{i=1}^{m} D_i \geq 0.$$

Values between 0.025 and 0.975 indicate that the model is adequate and cannot be rejected (*i.e.*, zero falls within the 95% posterior-predictive interval).

Posterior-predictive simulation requires: (1) the ability to simulate biogeographic data, given parameters of the model, and; (2) summary statistics that allow us to compare the resulting simulated datasets to the observed dataset. We describe each of these components below.

##### Simulation

Under a time-constant biogeographic model, we used the `sim.history()` function in the `R` package `phytools` (Revell 2012) to simulate full dispersal history forward in time over a phylogeny. We implemented a forward-simulation function in `R`, as an extension of the `sim.history()`, to perform the same simulation under a piecewise-constant geographic process. These functions allow us to perform posterior-predictive simulations to assess the adequacy of both the time-constant and the piecewise-constant models. The function we implemented is available in the `R` scripts included in our GitHub and Dryad repositories.

##### Summary statistics

We use two types of summary statistics: (1) statistics designed to assess global adequacy (*i.e.*, over the entire geographic history), and; (2) statistics designed to assess adequacy on an interval-by-interval basis.

To assess global model adequacy, we use two summary statistics: (1) the *parsimony statistic*, and; (2) the *tip-wise multinomial statistic*. For the parsimony statistic, we simply calculate the parsimony score for the given simulated or observed dataset using the `parsimony()` function in `R` package `phangorn` (Schliep 2010). The tip-wise multinomial statistic is similar to the multinomial statistic introduced by Goldman

(1993) and used in posterior-predictive simulation developed by Bollback (2002), which treats the sites (columns) in a molecular alignment as outcomes of a multinomial trial. Our tip-wise statistic is similar, but treats the states at the tips of the tree for the single geographic character (*i.e.*, site) as the outcomes of the multinomial trial. For the tip-wise multinomial statistic, we calculate:

$$T(G \mid \theta_i) = \sum_{i=1}^{k} n_i \ln(n_i/n),$$

where $n$ is the number of tips, and $n_i$ is the number of tips in state $i$. (Note that this statistic is also similar to the entropy statistic used to assess genetic variability along sequences; Shannon 1948; Schneider et al. 1986.)

To assess interval-specific adequacy, we use interval-specific variants of the two statistics described above. Specifically, these interval-specific summary statistics are calculated for each time interval to assess the adequacy of each model in describing the geographic dispersal pattern within that time interval. For the interval-specific parsimony statistic, we use the `ancestral.pars()` function in `R` package `phangorn` (Schliep 2010) to obtain the parsimonious ancestral-state reconstruction, and then compute the parsimony score in each interval for both the simulated and observed dataset. For the interval-specific tip-wise multinomial statistic, we simply calculate this statistic using the tips that were sampled in the focal interval for both the simulated and observed dataset. Details regarding the computation of these summary statistics are available in an `R` script, `posterior_predictive_teststatistics_functions.R`, included in our GitHub and Dryad repositories.
